# Population structure of *Bacillus cereus sensu lato* associated with foodborne outbreaks in France between 2004 and 2023

**DOI:** 10.1101/2025.03.24.644875

**Authors:** Ksenia Mozhaitseva, Sylvie Pairaud, Olivier Firmesse, Mathilde Bonis

## Abstract

*Bacillus cereus sensu lato* (*Bcsl*) is a group of closely related bacterial species known for their resistant spores, enabling them to persist in a dormant state and thereby colonize and adapt across diverse environments. *Bcsl* is known for its harmful impact on human health, producing toxins that cause emetic and diarrheal syndromes or provoking opportunistic infections in hospitals. Importantly, *Bcsl* is the most frequent confirmed or presumptive causative agent associated with foodborne outbreaks (FBOs) in France. In our study, we assessed the population structure of a large collection of *Bcsl* isolated during FBOs investigation in France between 2004 and 2023, focusing on the association between distinct populations and food categories. Using 294 genomes from 183 FBOs, we applied genomic clustering and phylogenomic analysis to accurately identify predominant *Bcsl* populations: *B. cereus sensu stricto* (17.0%), *B. paranthracis* (16.1%), and *B. thuringiensis* subsp. *kurstaki* (7.6%), which were positively associated with composite dishes, cereals, and vegetable-based salads, respectively. Some strains were phylogenetically closely related to clinical isolates, highlighting the need to assess the antibiotic susceptibility of *Bcsl*. Notably, one *Bcsl* clade, *B. cytotoxicus*, lacking beta-lactamase-encoding genes showed a greatly increased sensitivity to ampicillin than other *Bcsl* considered to be naturally resistant to beta-lactams. Additionally, some strains showed reduced susceptibility to macrolides and cyclins. Finally, we identified several signatures of recombination and horizontal transfer, suggesting the ability of *Bcsl* to share fitness-enhancing alleles and acquire virulence and antibiotic resistance genes by non-pathogenic strains. These findings would be valuable for improving the surveillance of *Bcsl*.

**HIGHLIGHTS:** *Bcsl* associated with French food poisonings consist of 14 clusters.

*Bcss,* the most frequent clade, is mainly found in composite dishes.

*Bcsl* toxin genes are present in various clades, suggesting horizontal gene transfer.

Some *Bcsl* linked to food poisonings or septicemia belong to the same populations.

*B. cytotoxicus,* lacking *bla* genes, showed increased sensitivity to ampicillin.

## 1. INTRODUCTION

*Bacillus cereus sensu lato* (*Bcsl*), or the *Bacillus cereus* group, comprises 28 validly and effectively published species (Parte et al., 2020). These bacteria are Gram-positive, aerobic or facultatively anaerobic, spore-forming, and primarily telluric and rhizospheric (Vilain et al., 2006). The spores of *Bcsl* are resistant to heat, UV radiation, acid, and desiccation, enabling them to persist in a dormant state and thereby contaminate diverse environments (Ehling-Schulz et al., 2019). In addition to soil, *Bcsl* has been isolated from plants, seawater, food products, animals, and humans (Schnepf et al., 1998; Guinebretière et al., 2013; Miller et al., 2016; Liu et al., 2017; Glasset et al., 2018; Tohya et al., 2021; Muigg et al., 2022; Makuwa et al., 2023; Robas Mora et al, 2023). *Bcsl* includes species known for their harmful impact on human health.

It is the most frequent confirmed or presumptive causative agent associated with foodborne outbreaks (FBOs) in France (Santé Publique France, 2024), and the second most frequently reported causative agent in Europe (European Food Safety Authority, 2024). *Bcsl* can produce various toxins, resulting in emetic or diarrheal syndromes. The emetic syndrome is caused by cereulide, a heat-stable, non-ribosomal cyclic peptide, whose synthesis is directed by the *ces* operon located in the pCER270 megaplasmid. In the most severe cases, ingestion of this toxin, produced in food, can lead to multiple organ failure and death (Rouzeau-Szynalski et al., 2020). Cereulide-producing strains are commonly referred to as emetic *B. cereus* and are predominantly found in specific *B. paranthracis* clades (Carroll and Wiedmann, 2020; Carroll et al., 2022b; Frentzel et al., 2024). However, they are also occasionally identified in other *Bcsl* (Mei et al., 2014; Pheepakpraw et al., 2023; Frentzel et al., 2024). The diarrheal syndrome is associated with the pore-forming enterotoxins: hemolysin BL (Hbl), non-hemolytic enterotoxin (Nhe), and cytotoxin K (CytK). Hbl and Nhe are tripartite toxins encoded by the *hblCDA* and *nheABC* operons, respectively (Tuipulotu et al., 2021). Cytotoxin K is encoded by the *cytK* gene existing in two variants: *cytK-1* and *cytK-2*. CytK-1 shows higher cytotoxicity and is restricted to *B. cytotoxicus* (Guinebretière et al., 2013), while CytK-2 can be produced by other *Bcsl* (Fagerlung et al., 2004). These enterotoxins, produced in digestive tract, damage intestinal epithelial cells by forming pores leading to osmotic lysis and, ultimately, diarrhea. Although less characterized, other virulence factors, such as sphingomyelinase, enterotoxin FM, hemolysins I, II and III, metalloproteases, and phospholipases complement the action of the *Bcsl* enterotoxins (Tuipulotu et al., 2021).

The members of the *B. cereus* group are also able to disseminate in hospital environments contaminating medical devices, equipment, and linen, and cause various local and systemic infections, particularly in immuno-compromised individuals (Glasset et al., 2018). Certain species, such as *B. hominis*, *B. sanguinis*, and *B. paramobilis*, were originally isolated from the blood of hospitalized patients (Tohya et al., 2021), and *B. basilensis* was first recovered from a wound swab under clinical conditions (Muigg et al., 2022). Moreover, in *B. mobilis*, originally isolated from ocean sediment (Liu et al., 2017), one strain was identified as clinically relevant and linked to polymicrobial catheter infection and enteritis (Muigg et al., 2022). This reinforces the need to properly define the antibiotic susceptibility pattern of *Bcsl.* To date, *Bcsl* has been described as naturally resistant to beta-lactams, while resistance to other families of antibiotics were described more sporadically (Glasset et al., 2018; Bianco et al., 2021; Jung et al., 2022; Rajalingam et al., 2022; Mohammadi et al., 2023).

Various *Bcsl* are known for their commercial use, in particular, *Bacillus thuringiensis* (*Bt*) which can produce delta-endotoxins (Cry, Cyt, and Vip) with insecticidal activity (Schnepf et al., 1998). Several *Bt* strains, primarily those belonging to subspecies *aizawai* (*Bta*) and *kurstaki* (*Btk*), have been commercialized and are widely used for pest control (Jallouli et al., 2020). Besides, certain strains of *B. toyonensis* are used as probiotics in animal feed (Pandith and Kumar, 2024). An isolate of another species, *B. mycoides,* is used as a fungicide for plant protection (Jacobsen et al., 2007).

Since the *B. cereus* group is of particular importance for FBO monitoring, it is crucial to properly assess the *Bcsl* population structure and link it to FBO data in order to identify the most prevalent and harmful *Bcsl* lineages. Each lineage could then be associated with multiple factors, such as specific food categories and epidemiological indicators.

However, in some cases, species assignment and proper definition of population boundaries within *Bcsl* present challenges due to its high genome plasticity (Ehling-Schulz et al., 2010; Böhm et al., 2015; Bazinet, 2017) and elevated genetic similarity between some species (Liu et al., 2015; Bazinet, 2017; Torres Manno et al., 2020). Historically, *Bcsl* members were classified into different species primarily based on phenotypic traits, such as colony morphology, growth temperature specificity, or association with symptoms and diseases (Tallent et al., 2020). Later, other criteria included presence of virulence plasmids, similarity measured using DNA-DNA hybridization techniques, phylogenetic groups and sequence types based on comparison of single locus and multiple loci (SLST and MLST), respectively (Caroll et al., 2022a). When applying the SLST approach, partial sequence of the gene encoding pantoate-β-alanine ligase (*panC*) allows a relatively high resolution to differentiate the major phylogenetic groups of *Bcsl,* I to VII (Guinebrètiere et al., 2008). To date, *panC* phylogenetic groups I, V, and VII are represented by *B. pseudomycoides*, *B. toyonensis* and *B. cytotoxicus*, respectively. Group IV includes *B. cereus sensu stricto* (*Bcss*) and *Bt*. Groups II, III, and VI are the most diverse and comprise several species (Caroll et al., 2020a). Among MLST schemes, the one based on seven housekeeping genes (Priest et al., 2004) is the most widely used. For example, using this scheme, certain sequence types (STs) were found to be frequently associated with foodborne outbreaks, such as emetic strains of *B. paranthracis* belonging to ST26 (Frentzel et al., 2024). The study of Tong et al. (2024) found that within *Bcss*, ST24 and ST142, both associated with foodborne illness, were the most prevalent STs in Europe.

Finer characterization of *Bcsl* is feasible thanks to whole genome sequencing data (Liu et al., 2015; Bazinet, 2017; Torres Manno et al., 2020). Currently, the assignment of *Bcsl* strains to genomospecies is based on the calculation of the Average Nucleotide Identity (ANI) index, the percentage of nucleotide identity in the alignment between query and reference (e.g. type strain) genomes. Commonly, an ANI equal to or higher than 95% reflects natural species boundaries (Jain et al., 2018). However, this 95% threshold is inadequate for the *B. cereus* group, leading to ambiguous genomospecies assignment results, as discussed in Caroll et al. (2020b, 2021). To avoid overlapping genomospecies within *Bcsl*, the authors proposed an ANI threshold of 92.5%. Using this threshold, both *Bcss* and *Bt* are grouped into the same *Bcss* genomospecies, while all species belonging to *panC* groups II (except for *B. luti*) and III are grouped into the *B. mosaicus* genomospecies. Any strain carrying a plasmid that harbors Cry/Cyt/Vip toxin, emetic toxin and/or anthrax toxin encoding genes is labeled with the corresponding biovar name (Thuringiensis, Emeticus, and/or Anthracis, respectively), appended to the genomospecies name. This nomenclature is useful for clinicians as it helps avoid ambiguous genomospecies assignment and enables the rapid identification of genotypes with clinical and industrial relevance. However, it is less suited for finer resolution, particularly for *panC* groups III and IV. *Bt*, widely used as a biopesticide, can be confused with the phylogenetically closely related *Bcss,* a known food poisoning agent. The latter may carry the plasmids harboring insecticidal toxin genes acquired via horizontal transfer (Battisti et al., 1985). Furthermore, these two species show similar phenotypic characteristics (van Tongeren et al., 2014), highly similar chromosomal structure (Carlson et al., 1996), and an ANI greater than 96% (Torres Mano et al., 2020). Therefore, it is of particular importance to accurately distinguish *Bt* from *Bcss*, given increasing data on the *Bt* presence in food products, its association with foodborne outbreaks (Frentzel et al., 2020; Bonis et al., 2021; Biggel et al., 2022a, 2022b) and nosocomial infections (Kuroki et al., 2009; Bianco et al., 2021; Barakat et al., 2024).

*Bcsl* is characterized by a dynamic pangenome due to its high genome plasticity (Ehling-Schulz et al., 2010; Böhm et al., 2015; Bazinet, 2017). The pangenome of *Bcsl* includes a core genome, which contains genes shared by 95–100% of strains within a bacterial species, and an accessory genome, which consists of genes found in only some strains. Using 146 *Bcsl* genome assemblies, Bazinet (2017) estimated its pangenome at ∼60,000 genes, while the core genome at ∼600 genes, thereby suggesting high adaptability of *Bcsl* resulting from the acquisition of accessory genes. *Bcsl* accessory genes are enriched in various functions including those linked to antimicrobial resistance and virulence (White et al., 2022). Furthermore, Böhm et al. (2015) demonstrated that toxin-encoding accessory genes, such as *hbl* and *cytK*, were horizontally transferred between *Bcsl* clades. The authors concluded that this could enable non-pathogenic strains to acquire virulence factors. This might have consequences for risk assessment of *Bcsl*.

The aim of this study was to resolve the population structure of *Bcsl* isolated from foodborne outbreaks in France between 2004 and 2023, with a focus on the association between predominant populations and various food categories. Using a large collection, consisting of 294 *Bcsl* genomes from 183 FBOs, we applied genome clustering method followed by phylogenomic analysis to accurately determine the predominant FBO-associated *Bcsl* populations. We identified (i) 11 *Bcsl* clusters composed of single species and (ii) three composite clusters composed of closely related species, each consisting of several populations. We revealed three predominant populations: a single *B. cereus sensu stricto* population, one *B. paranthracis* population including emetic strains, and *B. thuringiensis* subsp. *kurstaki*, which prevailed in composite dishes, cereals, and vegetable-based salads, respectively. In addition, several FBO-associated *Bcsl* strains belonged to the same population as clinical strains, highlighting the need to assess the antibiotic susceptibility of *Bcsl.* We also found that one population, *B. cytotoxicus*, was significantly more sensitive to ampicillin, than the other populations. Finally, we revealed several signatures of homologous recombination and horizontal transfer, which enable non-pathogenic strains to acquire fitness-enhancing genetic variants, as well as virulence and antibiotic resistance genes. These findings should be considered to improve the surveillance of *Bcsl*.

## 2. MATERIALS AND METHODS

### 2.1. Genome assemblies

We analyzed a collection of 294 draft genomes of *Bcsl* (Supplementary Table S1) sampled during foodborne outbreaks in France between 2004 and 2023 and stored in the Food Safety Laboratory strain bank of the French Agency for Food, Environmental and Occupational Health and Safety (ANSES). The corresponding raw reads are available at the NCBI platform under the projects PRJNA1209675, PRJNA781790, and PRJNA547495. The DNA extraction technique and the sequencing protocol are described in Bonis et al. (2021). The genomes were assembled using the in-house *bacflow* pipeline, as described in Fichant et al. (2024). The assembly size varied from 4.1 to 7.0 Mbp, the N50 statistic ranged from 15.9 to 936 kbp, and the L50 values ranged from 2 to 91. The assembled genomes of 23 clinical and blood culture isolates were retrieved from the MicroBIGG-E browser (https://www.ncbi.nlm.nih.gov/pathogens/microbigge/, accessed in June 2024). The data were filtered by location (France) and isolation source (blood, patient, intravascular catheter) (Supplementary Table S2).

### 2.2. Species tree

An unrooted phylogenetic tree was constructed based on 1723 protein sequences of single-copy orthologous genes identified across the *Bcsl* genomes using OrthoFinder v.2.5.5 (Emms and Kelly, 2015). Multiple protein sequence alignment was performed by MUSCLE v.3.8.1551 (Edgar, 2004). The resulting ‘SpeciesTreeAlignment.fa’ file was used to infer a maximum likelihood phylogeny in IQ-TREE v.2.2.6 (Minh et al., 2020). The best protein substitution model identified by ModelFinder (Kalyaanamoorthy et al., 2017) (-m MFP tag in IQ-TREE) was JTT+F+I+R10 (Jones et al., 1992; Yang, 1995; Soubrier et al., 2012). The number of replicates for ultrafast bootstrap approximation (Hoang et al., 2018) for the branch support assessment was set to 1000 (-B 1000 tag in IQ-TREE).

### 2.3. Taxonomic assignment, *panC* group, and *Bt* typing

Taxonomic assignment of *Bcsl* genomes was performed using the framework of Caroll et al. (2020b) implemented in Btyper3 v.3.4.0 (Caroll et al., 2020a). For each genome, the tool identified the genomospecies, the closest type strain, and the PubMLST (https://pubmlst.org) sequence type. The samples were further assigned to one of the *panC* (I to VII) phylogenetic groups proposed by Guinebrètiere et al. (2008) using a publicly available tool (https://toolcereusid.shinyapps.io/Bcereus/; accessed in March 2024). In addition, *Bta* and *Bt*k were identified based on the presence of *Bt-*specific genes following the workflow described in Fichant et al. (2022) and implemented in Bt_typing (https://github.com/afelten-Anses/Bt_typing; accessed in March 2024).

### 2.4. Genome clustering

The clustering of *Bcsl* genomes was performed using PopCOGenT (Arevalo et al., 2019) (https://github.com/philarevalo/PopCOGenT, accessed February 2024), designed to identify bacterial populations by calculating a distribution of identical DNA sequence lengths between pairs of genomes. PopCOGenT first reconstructs networks (clusters) of similar genomes and further resolves subclusters representing putative populations.

### 2.5. Pangenome examination

The *Bcsl* genomes were annotated with Bakta v.1.9.3 (Schwengers et al., 2021) using the Bakta database v.5.1 (January 2024). The resulting *.gff3* annotations were processed with Panaroo v.1.4.1 (Tonkin-Hill et al., 2020) with default parameters. The pangenome was reconstructed at two levels: (i) across all members of *Bcsl* and (ii) within a specific PopCOGenT cluster of interest. Further hierarchical clustering of the *Bcsl* genomes was based on gene presence-absence and performed using the gene_presence_absence.Rtab output matrix from Panaro. Pairwise Jaccard distances were calculated by applying the dist (method=”binary”) R function. The resulting distance matrix was used in the pvclust (method.hclust=”complete”, nboot=1000) package v.2.2.0 (Suzuki et al., 2019) implemented in R v.4.3.2 (R Core Team, 2023). The genomes sharing similar accessory gene content were clustered into the Groups. The Group was considered significant at approximately unbiased *p*-value >95% and bootstrap probability >95%. Thereafter, candidate Group-specific genes were identified by Scoary v.1.6.16 (Brynildsrud et al., 2016) with thresholds of 80% sensitivity, 80% specificity, and a *p*-value cutoff of 0.05 corrected for multiple testing by Benjamini-Hochberg method. Further Cluster of Orthologous Groups (COG) enrichment analysis was performed in MicrobiomeProfiler (Wu et al., 2021) at two levels: (i) PopCOGenT cluster against the entire background *Bcsl* COG set, and (ii) Groups found within the PopCOGent cluster of interest against the entire background cluster COG set. Significant COGs were identified at an adjusted *p*-value cutoff set to 0.05 (Benjamini-Hochberg correction).

### 2.6. Population structure

We carried out a population structure analysis for each composite PopCOGenT cluster consisting of several subclusters. First, we applied fastGEAR (Mostowy et al., 2017) which uses hidden Markov model to assess shared ancestry and reveal putative recombining regions within homologous regions of bacterial genomes. The tool defines lineages as groups of samples that share identical sequences at least 50% of the sites. Recombination events were inferred by identifying regions with a log Bayes factor (logBF)>0 indicating a higher probability of recombination than mutation. We focused on both recent and ancestral recombination. Ancestral recombination was defined by fastGEAR as putative recombination event affecting all samples in the lineage, while recent recombination referred to events occurring in some but not all isolates. As input, we used a filtered alignment of a set of core genes reconstructed by Panaroo v.1.4.1 (Tonkin-Hill et al., 2020) for each PopGOCenT cluster. Ambiguous sites and indels were excluded from the alignment using snp-sites v.2.5.1 (Page et al., 2016). Second, we validated results by assessing recombination with ClonalFrameML (Didelot and Wilson, 2015) which employs a maximum likelihood method. The program estimates an average frequency of recombination over mutation (R/θ), an average number of substitutions (δν) per recombination event, and an overall impact of recombination on phylogeny (r/m=R/θ×δν) in a population of interest. A guide phylogenetic tree was reconstructed for each cluster with IQ-TREE v.2.2.6 (Minh et al., 2020) using the filtered alignment of core genes, the best nucleotide substitution model identified by ModelFinder (Kalyaanamoorthy et al., 2017), and 1000 replicates for ultrafast bootstrap approximation (Hoang et al., 2018). Thereafter, the tree topology was refined by excluding regions affected by recombination

### 2.7. Identification of toxin, antimicrobial and heavy metal resistance genes

The *cesABCD, nheABC* and *hblCDA* operons, *cytK-1* and *cytK-2* genes, and the genes encoding insecticidal toxins (Cry, Cyt, and Vip) were detected using Btyper3 v. 3.4.0 (Caroll et al., 2020a) with default parameters. The *hlyII* gene was directly searched in the genome annotations. Antimicrobial and heavy metal resistance genes were identified by AMRFinderPlus v.3.12.8 applying default parameters and using the reference protein database v.2024-01-31.1 (Feldgarden et al., 2021).

### 2.8. Antibiotic susceptibility testing

Antibiotic susceptibility of the *Bcsl* isolates was assessed using the Kirby–Bauer disk diffusion method in accordance with the European Committee on Antimicrobial Susceptibility Testing (EUCAST) methodology (method v13.0 and breakpoint tables for interpretation v15.0; https://www.eucast.org; accessed in January, 2025). Briefly, after isolation on trypticase soy agar-yeast extract (TSA-YE, bioMérieux) medium for 18h at 30°C, bacteria were resuspended in peptone salt broth (bioMérieux) at 0.5 McFarland turbidity and inoculated onto Mueller Hinton agar (Condalab) plates. Antibiotic disks (Oxoid) containing ampicillin (10µg, AMP10), erythromycin (15µg, E15), tetracyclin (30µg, TE30) and vancomycin (5µg, VA5) were then applied onto inoculates. Plates were incubated for 18h at 35°C before measuring the inhibition diameters.

### 2.9. Proportions of FBO-related *Bcsl* populations

To minimize the risk that the set of 294 available sequenced samples might not accurately reflect the true proportions of the FBO-associated *Bcsl* populations, we filtered the dataset as follows. First, we reviewed all ANSES FBO reports (n=560) indicating the presence of *Bcsl,* whether sequenced or not. The reports provided information on (i) the *panC* phylogenetic group determined by PCR, and (ii) the microscopic observation of insecticidal toxin crystals (*Bt* phenotype). Second, based on the reports, we calculated the proportion of each *panC* group: I – 0%, II – 15%, III – 38%, IV – 35%, V – 5%, VI – 5%, and VII – 2%. Within *panC* group IV, we identified the *Bt* phenotype based on its ability to produce insecticidal crystals. The *Bcss*:*Bt* ratio was estimated at 2:1. Third, we applied the bootstrap technique by generating 100 random sample sets in R v.4.3.2 (R Core Team, 2023) with respect to the obtained proportions, allowing no more than 1% of deviation. In each random sample set, the maximum possible number of samples, while respecting necessary proportions, was 147. Finally, from the generated sets, we retrieved the average percentage for each *Bcsl* population.

### 2.10. Association of *Bcsl* populations with food categories

After estimating proportions of the *Bcsl* populations, we then examined their distribution across various food categories. Information on contaminated products (Supplementary Table S1) was available in the FBO reports. Food items were classified using both the FoodEx2 (exposure hierarchy: Level 1 and 2) (European Food Safety Authority, 2015) and Food and Agriculture Organization (FAO) (Level 1 and 2) (Leclercq et al., 2019) classifications. Mean percentages of the *Bcsl* populations were calculated for each food category, based on the previously simulated datasets (see above). To determine whether a specific *Bcsl* population was predominantly associated with a particular food category, we performed a Fisher’s exact test (fisher.test function) using R v.4.3.2 (R Core Team, 2023). For each food category, we constructed a contingency table comparing mean counts of the *Bcsl* population of interest with the mean counts of the entire *Bcsl* group (background). Associations were considered significant at *p*<0.05.

### 2.11. Data visualization

Phylogenetic trees and plots were designed using ggtree v.3.5.1 (Yu et al., 2017; Xu et al., 2021; Yu, 2022), ape v.5.8 (Paradis and Schliep, 2019) and ggplot2 v.3.10.1 (Wickham, 2016) packages implemented in R v.4.3.2 (R Core Team, 2023).

## 3. RESULTS

### 3.1. *Bcsl* phylogeny and clustering

The studied *Bcsl* isolates were assigned to seven *panC* phylogenetic groups and divided into 14 PopCOGenT (0 to 13) clusters and 12 unclustered samples (Fig. 1, Table 1, Supplementary Table S1). A genome was considered unclustered if it represented a single sample within the cluster, assigned to a specific *Bcsl* species. For eleven clusters, PopCOGenT did not suggest any division into subclusters. We identified cluster 2 (n=15, *B. cytotoxicu*s), cluster 3 (n=5, *B. mobilis*), cluster 6 (n=11, *B. mycoides*), cluster 7 (n=3, *B. luti*), cluster 8 (n=9, *B. toyonensis*), cluster 12 (n=2, *B. paramobilis*), and cluster 13 (n=2, *B. wiedmannii*). Both cluster 4 (n=11) and cluster 11 (n=2) showed the highest ANI with the *B. anthracis* type strain, with respective ANI 97.2 to 98.7 and 94.3. The samples from both clusters did not possess the *pagA*, *lef*, and *cya* anthrax-associated genes. Two other clusters, 9 (n=8, *Bt*) and 10 (n=2, *Bt*), showed the highest ANI with the *Bt* Berliner type strain: 96.0-96.4 and 97.3-98.3 ANI, respectively. By contrast, three PopCOGenT clusters demonstrated composite structure and contained several subclusters (populations). Cluster 0 (n=75) consisted of nine PopCOGenT subclusters (0.0 to 0.8) containing samples exhibited the highest ANI with either the *B. paranthracis*, *B. pacificus* or *B. tropicus* type strains. Therefore, the cluster was named the *B. paranthracis* + *B. pacificus* + *B. tropicus* cluster. Another composite cluster, cluster 1 (n=113), comprised samples closely related to the *Bcss* Frankland and Frankland type strain and divided into subclusters 1.0 and 1.1. The cluster was named *Bcss* + *Bt* as additional analysis revealed that one of the subclusters included several *Bt* strains (see below). Finally, cluster 5 (n=24) comprised samples closely related to either the *B. wiedmanni* or *“B. pretiosus”* type strains and therefore named the *B. wiedmanni* + *“B. pretiosus”* cluster. The cluster was divided into five subclusters, 5.0 to 5.4. Subsequently, finer population structure analysis was performed on these three composite clusters.

**Figure 1.**
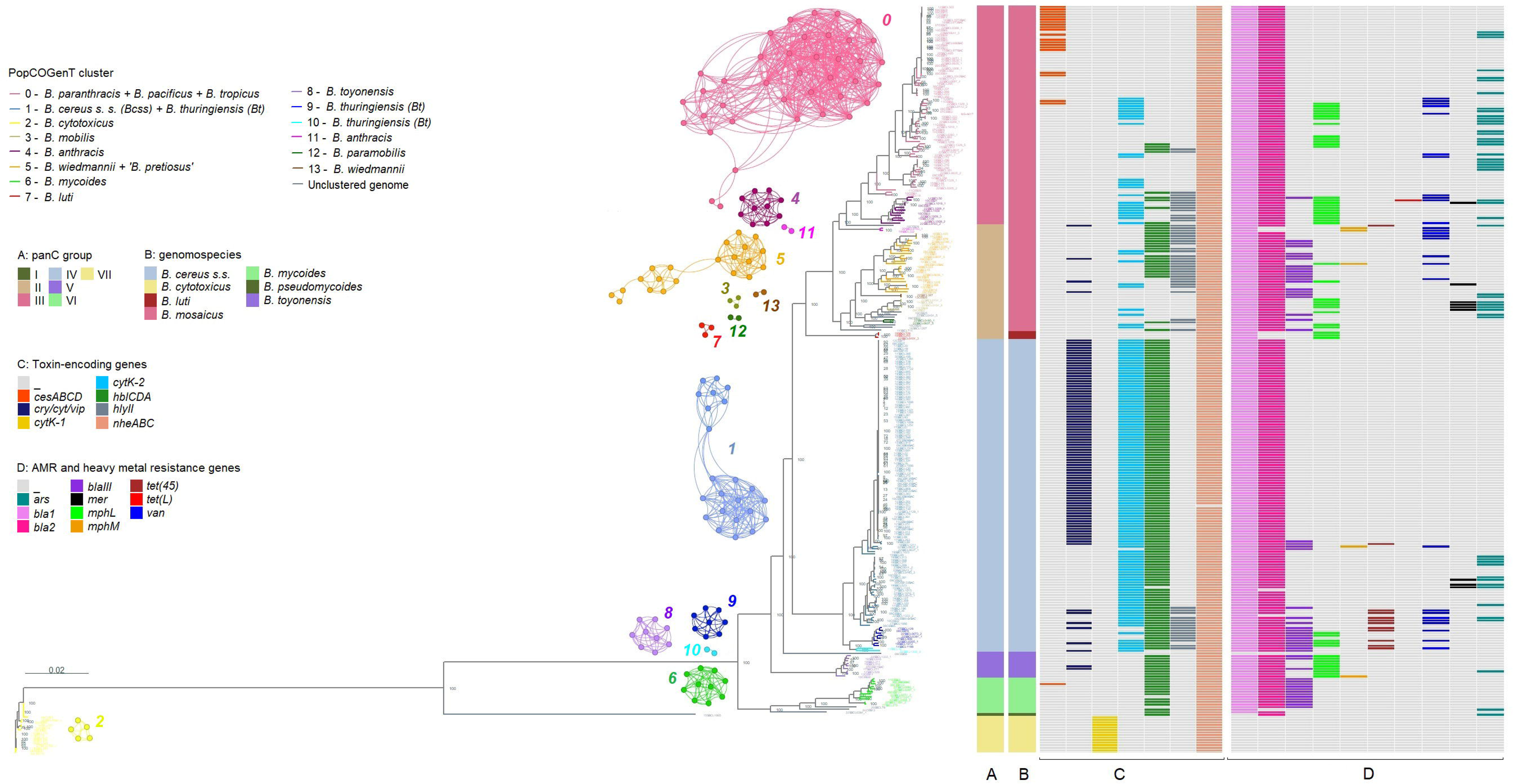
Maximum likelihood phylogenetic tree of *Bcsl* reconstructed using single-copy orthologous protein sequences. Branch lengths are presented in units of the expected number of amino acid substitutions per site. Bootstrap values are shown on internal nodes. Each sample label is colored according to the PopCOGenT cluster and corresponds to networks designed in Gephi v. 0.10.1. Each node corresponds to one sample or clonal complex, i.e. genomes showing high similarity (Arevalo et al., 2019). **A**: *panC* phylogenetic groups. **B**: genomospecies identified using the framework of Caroll et al. (2020a, 2020b). **C**: Toxin-encoding genes and operons: emetic toxin (*cesABCD*), insecticidal toxins (*cry/cyt/vip*), cytotoxin K-1 (*cytK-1*), cytotoxin K-2 (*cytK-2*), hemolysin BL (*hblCDA),* hemolysin II (*hlyII*), non-hemolytic enterotoxin (*nheABC*). **D**: Antimicrobial and heavy metal resistance genes: arsenic resistance operon (*ars*), beta-lactamase (*bla1, bla2, blaIII*), mercury resistance operon (*mer*), erythromycin resistance (*mphL*, *mphM*), tetracycline resistance (*tet(45), tet(L)*), regulatory part of the *van* operon (*vanRS*).

**Table 1.**
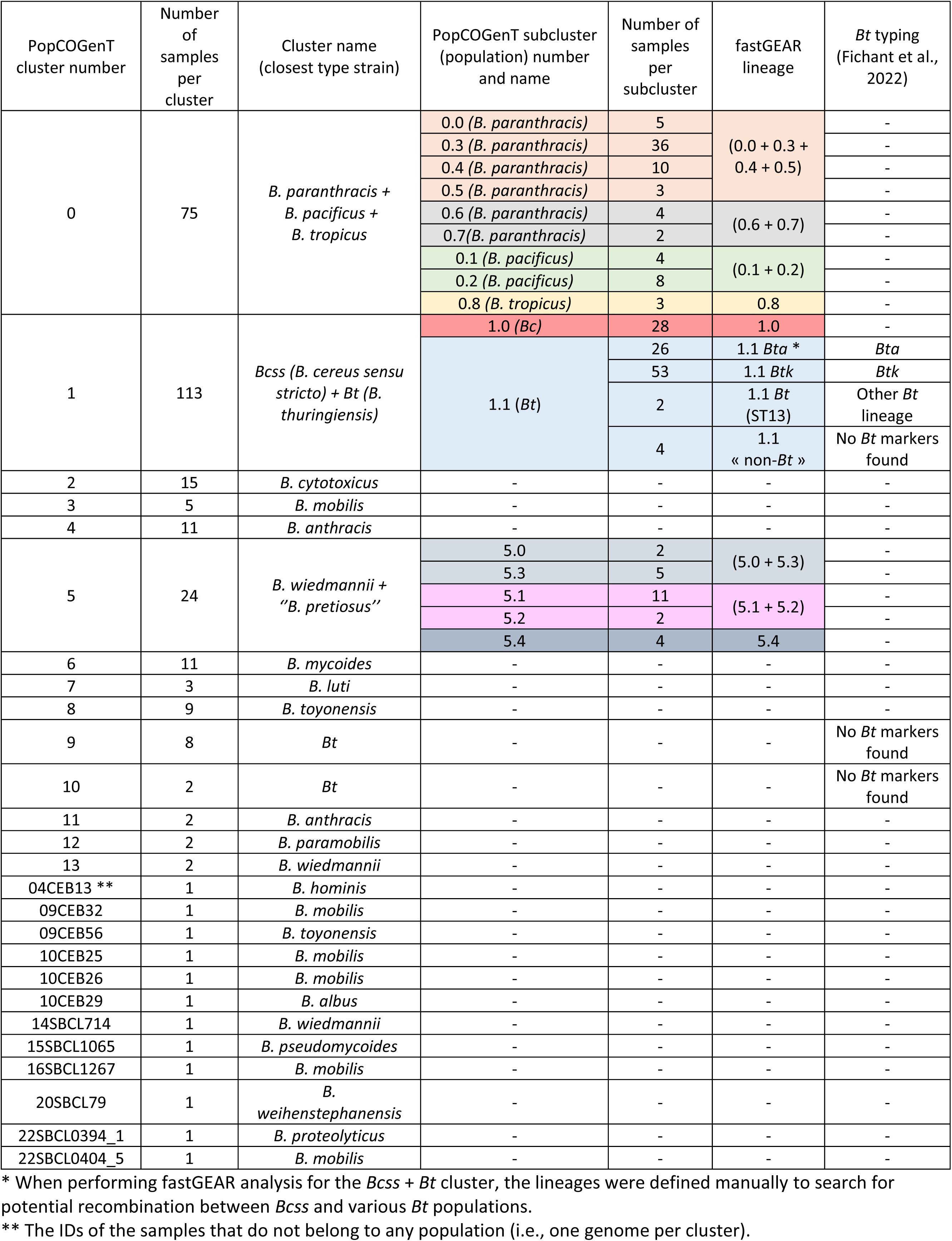
Clustering of *Bacillus cereus sensu lato* associated with foodborne outbreaks in France.

### 3.2. Population structure of B. paranthracis + B. pacificus + B. tropicus cluster 0

The first composite PopCOGenT cluster, cluster 0, consisted of nine distinct subclusters, 0.0 to 0.8 (Fig. 2A, Table 1). Six of these subclusters (0.0, 0.3, 0.4, 0.5, 0.6, and 0.7) were closely related in terms of ANI to the *B. paranthracis* type strain. Among them, subcluster 0.3 was the largest and included ST26, the most dominant sequence type. Subclusters 0.1 and 0.2 represented two distinct *B. pacificus* populations, while subcluster 0.8 corresponded to a *B. tropicus* population. Additional phylogenetic analysis revealed that the various clinical *B. paranthracis* samples did not form a distinct population. These clinical samples were phylogenetically close to either subclusters 0.3, 0.4, or 0.5. The most prevalent ST within the clinical samples was ST26 (11 out of 16 samples). Subsequently, fastGEAR detected lineages and admixture blocks indicating either shared ancestry or putative recombination (Fig. 2B, Fig. 2C, Table 1). The first fastGEAR lineage grouped the PopCOGenT subclusters 0.0, 0.3, 0.4 and 0.5. The second lineage consisted of subclusters 0.6 and 0.7. The third lineage consisted of subclusters 0.1 and 0.2. The fourth lineage corresponded to subcluster 0.8. Next, fastGEAR assessed putative recent recombination events. The highest total length of recombining regions was detected in the sample 18SBCL174 (subcluster 0.7, 1.11 Mbp), followed by 11CEB28 (0.8; 936.1 kbp), 22SBCL0891_1 (0.7; 831.6 kbp), and 19SBCL1079 (0.4; 456.6 kbp) (Fig. 2C). However, further analysis with ClonalFrameML showed the sample 11CEB28 (subcluster 0.8) to exhibit the highest sum length affected by recombination, estimated at 91.5 kbp, followed by 14SBCL19 (0.8; 30.2 kbp), 22SBCL1010_3 (0.6; 26.9 kbp), 22SBCL0891_1 (0.7; 25.8 kbp), and 18SBCL174 (0.7; 24.8 kbp) (Fig. 2A). For the entire cluster, the R/θ ratio was estimated at 0.75, indicating recombination to occur about 1.3 times less than mutation. However, the average number of substitutions introduced per recombination event (δν) was 7.7. Thus, the r/m parameter was 5.8. This indicates that, in this cluster, the recombination process overall led to 5.8 times more substitutions than the mutation process.

**Figure 2.**
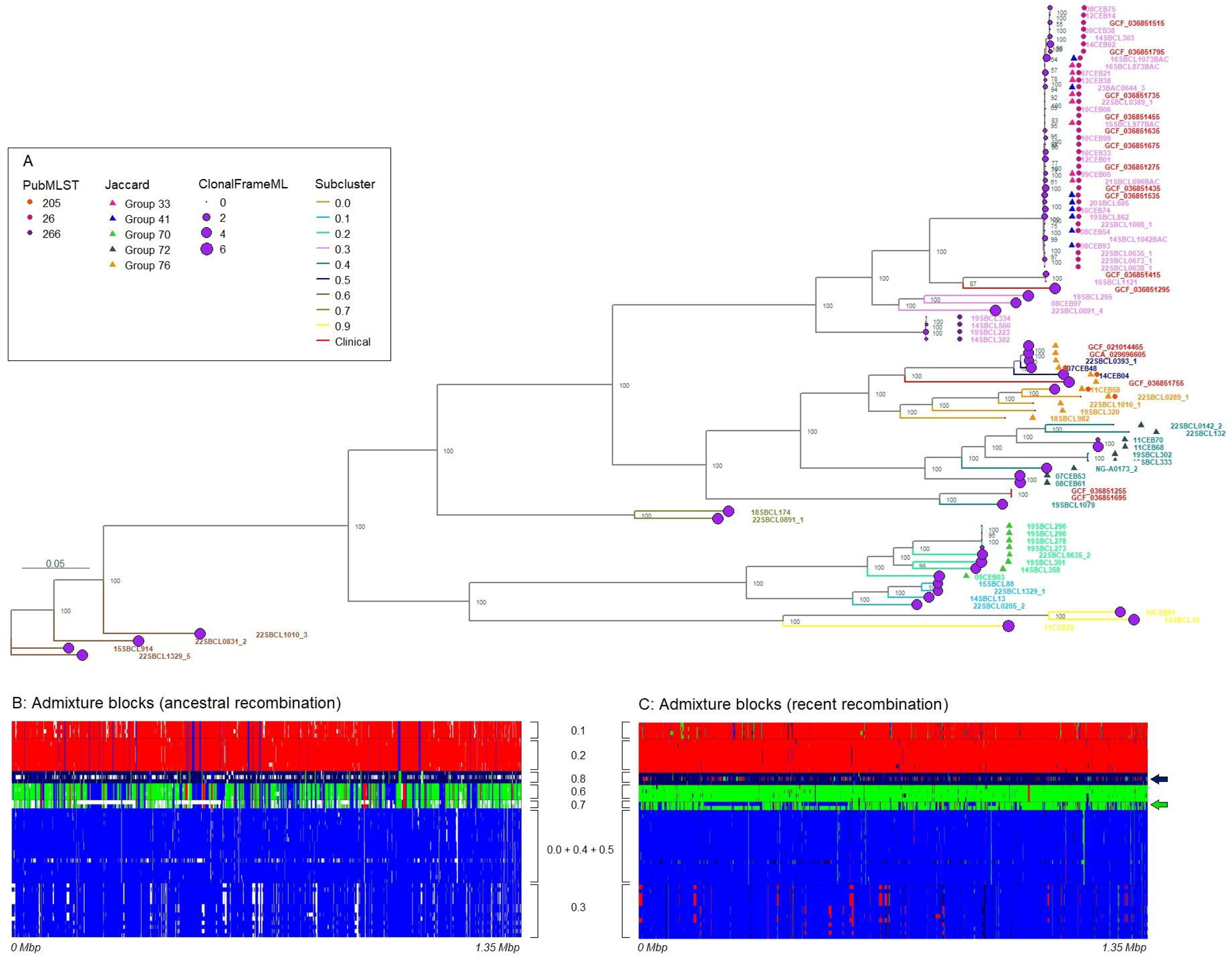
Maximum likelihood phylogenetic tree of the *B. paranthracis + B. pacificus + B. tropicus* PopCOGenT cluster 0. The tree was reconstructed using the alignment of core genes and the GTR+F+I+R5 substitution model (Tavaré, 1986). Branch lengths are presented in units of the expected number of nucleotide substitutions per site. Bootstrap values are shown at internal nodes. Each sample label is colored according to the corresponding PopCOGenT subcluster. **A**: PubMLST, predominant sequence types. Jaccard, Groups of genomes with similar accessory gene content. ClonaFrameML, logarithmic sum of recombining regions in the alignment of core genes; Subcluster, PopCOGenT subcluster number. **B**: Admixture blocks (ancestral recombination), patterns of shared ancestry within homologous regions across the core genome alignment. Blocks of the same color represent homologous regions shared by two lineages. The color of each block is assigned based on the lineage with the higher number of samples. FastGEAR detected four lineages: red (*B. pacificus* subclusters 0.1 + 0.2), dark blue (*B. tropicus* subcluster 0.8), green (*B. paranthracis* subclusters 0.6 + 0.7), and blue (*B. paranthracis* subclusters 0.0 + 0.3 + 0.4 + 0.5). White blocks represent recent recombination, indicating putative recombination events occurring in some, but not all, members of a population. **C**: Admixture blocks (recent recombination). The dark blue arrow indicates the sample 11CEB28. The green arrow indicates the sample 18SBCL174. Black blocks indicate putative external recombination. The displayed core genome alignment covers 1.35 Mbp out of the total 2.70 Mbp.

Gene content clustering revealed that the samples from *B. pacificus* subcluster 0.2 shared a similar set of accessory genes (Group 70) (Fig. 2A, Supplementary Fig. S1). Likewise, the analysis revealed a similar accessory genome within *B. paranthracis* subcluster 0.4 (Group 72). Notably, Group 76, which included genomes from *B. paranthracis* subclusters 0.0 and 0.5, was enriched in accessory genes associated with the ‘Cell wall/membrane/envelope biogenesis’ function (M COG) (Supplementary Table S3). Additionally, within *B. paranthracis* ST26 (subcluster 0.3), we identified two distinct groups, Group 33 and Group 41, each characterized by its own unique set of accessory genes.

### 3.3. Population structure of *Bcss* + *Bt* cluster 1

The second composite PopCOGenT cluster, which included the highest number of samples (n=113), composed of two subclusters: *Bcss* subcluster 1.0 and *Bt* subcluster 1.1 (Fig. 3A, Table 1). An additional application of the workflow to identify *Bt*-specific markers (Fichant et al., 2022) revealed a finer structure within the *Bt* subcluster. This analysis identified *Bt* subsp. *aizawai* (*Bta*, ST15), *Bt* subsp. *kurstaki* (*Btk*, ST8), two *Bt* ST13 samples (1.1 *Bt*), and four genomes in which no *Bt*-specific markers described in Fichant et al. (2022) were detected (designated hereafter as 1.1 *“*non-*Bt”*). Further fastGEAR analysis revealed admixture blocks corresponding to putative recent recombination in the 1.1 *“*non-*Bt”* genomes and the *Bcss* subcluster, but not in *Bta*, *Btk*, or 1.1 *Bt* ST13 (Fig. 3B). Admixture blocks corresponding to ancestral recombination were not detected. ClonalFrameML similarly detected recombination in the 1.1 *“*non-*Bt”* samples and in the *Bcss* subcluster 1.0. The maximum length of recombining regions was detected for the sample 15SBCL85 (1.1 *“*non-*Bt”*; 46.8 kbp), followed by 08CEB85 (*Bcss*; 21.9 kbp), 19SBCL1023 (1.1 *“*non-*Bt”*; 14.5 kbp) and 17SBCL1086 (*Bcss*; 14.5 kbp). For the entire *Bcss + Bt* cluster 1, the average R/θ, δν and r/m parameters were estimated at 0.59, 7.3 and 4.3, respectively. Additionally, the reconstructed phylogeny revealed that seven clinical *Bcss* samples did not form a distinct population but were phylogenetically closely related to various FBO-related strains (Fig. 3A).

**Figure 3.**
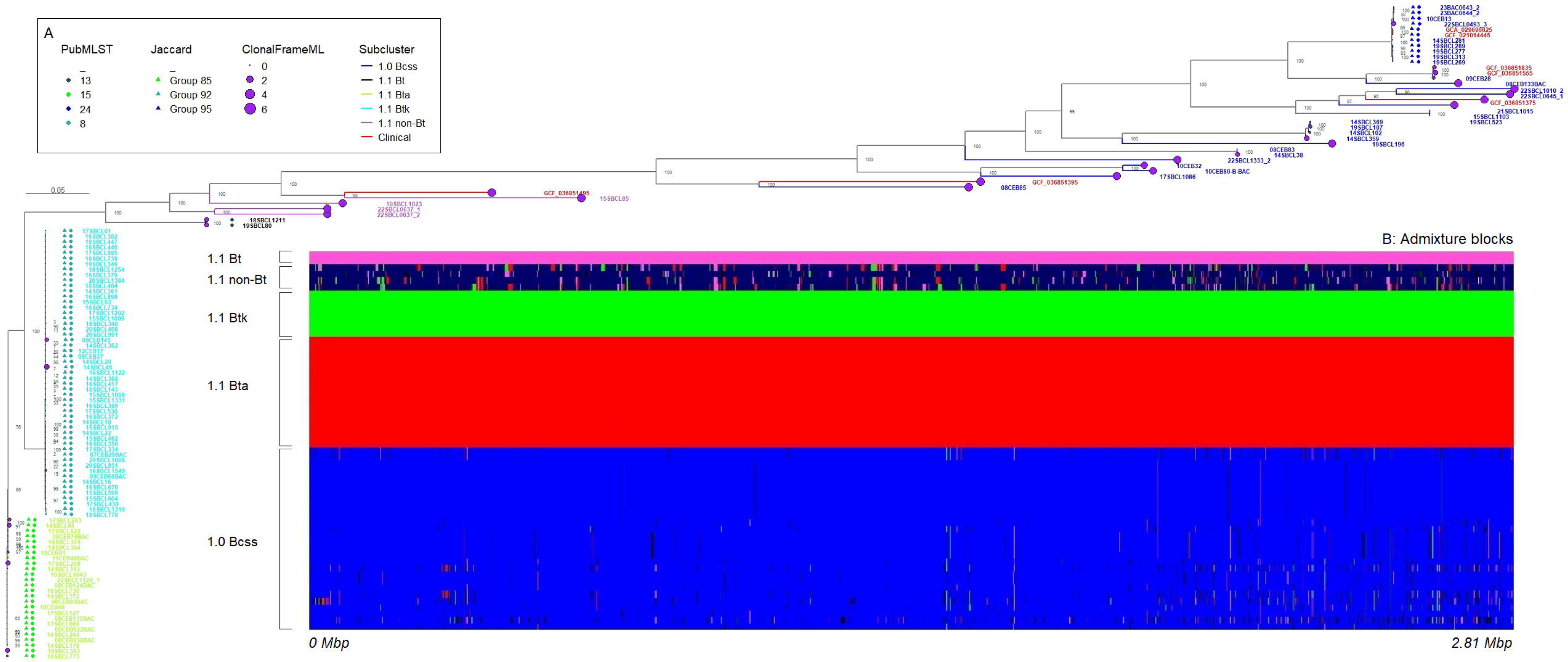
Maximum likelihood phylogenetic tree of the *Bcss* and *Bt* PopCOGenT cluster 1. The tree was reconstructed using the alignment of core genes and the GTR+F+I+R3 substitution model (Tavaré, 1986). Branch lengths are presented in units of the expected number of nucleotide substitutions per site. Bootstrap values are shown at internal nodes. Each sample label is colored according to the corresponding PopCOGenT subcluster. **A**: PubMLST, predominant sequence types. Jaccard, groups of genomes with similar accessory gene content. ClonaFrameML, logarithmic sum of recombining regions in the alignment of core genes. Subcluster, PopCOGenT subcluster number. **B**: Admixture blocks (ancestral recombination), patterns of shared ancestry within homologous regions across the core genome alignment. Blocks of the same color represent homologous regions shared by two lineages. The color of each block is assigned based on the lineage with the higher number of samples. FastGEAR detected five lineages: purple (*Bt*), dark blue (“non-*Bt*”), green (*Btk*), red (*Bta*), and blue (*Bcss*). Black blocks indicate putative external recombination.

Gene content analysis revealed several groups of genomes with similar accessory genes. Within the *Bt* subcluster 1.1, we identified the *Bta* (Group 85) and *Btk* (Group 92) significant groups (Fig. 3A, Supplementary Fig. S2). Within the *Bcss* subcluster, the largest significant group (Group 95) comprised 11 samples corresponding to *Bcss* ST24. Interestingly, the 1.1 *“*non-*Bt”* samples showed greater similarity to *Bcss* than to *Bta* or *Btk* in terms of gene content. Furthermore, enrichment analysis performed for the significant Groups revealed that *Bta* and *Bcss* ST24 each possessed a specific set of accessory genes enriched in the ‘Mobilome: prophages, transposons’ functional category (X COG) (Supplementary Table S3).

### 3.4. Population structure of *B. wiedemanni + “B. pretiosus”* cluster 5

The third analyzed composite cluster, cluster 5, comprised two species, *B. wiedemanni* and *“B. pretiosus”*. Within this cluster, PopCOGenT identified five distinct subclusters (populations), 5.0 to 5.4 (Fig. 4A, Table 1). Admixture analysis, performed by fastGEAR, identified three distinct lineages (Fig. 4B, Table 1). The first lineage grouped subclusters 5.0 and 5.3, the second grouped subclusters 5.1 and 5.2, and the third corresponded to subcluster 5.4. In the core genome alignment (Fig. 4B), we identified several admixture blocks shared between lineages (5.0 + 5.3) and 5.4, spanning from 0.509 to 0.750 Mbp and 1.403 to 1.685 Mbp. Additionally, other admixture regions (0.081 to 0.114 Mbp, 0.658 to 0.828 Mbp, and 1.033 to 1.073 Mbp) were shared between the (5.1 + 5.2) and (5.0 + 5.3) fastGEAR lineages. Recombination analysis performed by ClonalFrameML, revealed the highest sum of recombining regions in the sample 11CEB61 (0.77 Mbp), followed by 17SBCL1228 (0.61 Mbp), 08CEB128 (0.58 Mbp), 08CEB132BAC (0.55 Mbp), and 09CEB40 (0.52 Mbp) (Fig. 4A). The R/θ, δν, and r/m parameters were estimated at 0.27, 8.9, and 2.4, respectively. The analysis of accessory gene content identified significant groups that did not follow the population structure (Fig. 4A, Supplementary Fig. S3). Group 12 included samples from subclusters 5.0 and 5.3. Group 13 consisted of three genomes from subcluster 5.4, along with the samples 14SBCL108 and 17SBCL1228 that belonged to subclusters 5.1 and 5.3, respectively.

**Figure 4.**
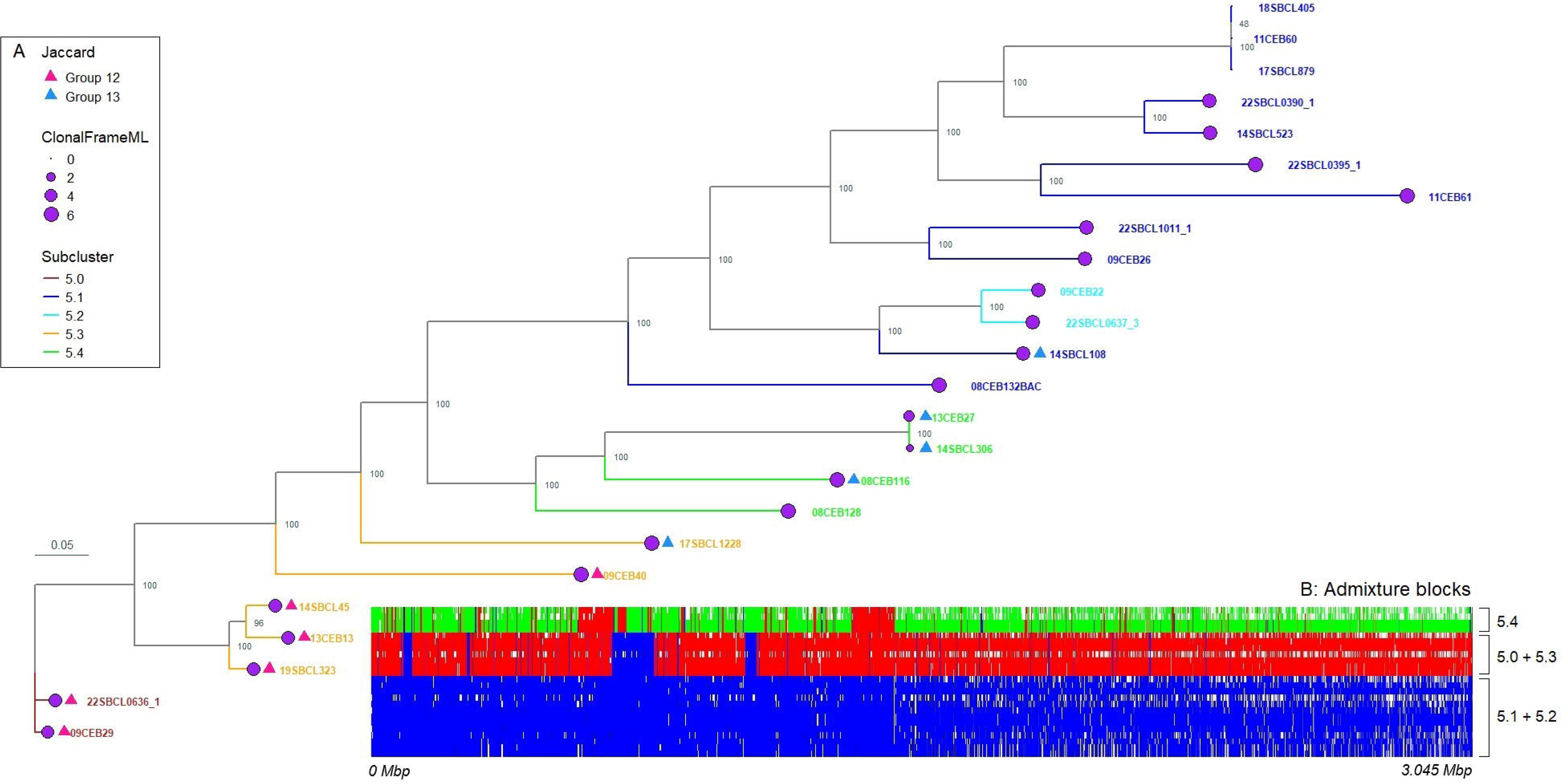
Maximum likelihood phylogenetic tree of *B. wiedmannii* and “*B. pretiosus*” PopCOGenT cluster 5. The tree was reconstructed using the alignment of core genes and the GTR+F+I+R7 substitution model (Tavaré, 1986). Branch lengths are presented in units of the expected number of nucleotide substitutions per site. Bootstrap values are shown at internal nodes. Each sample label is colored according to the corresponding PopCOGenT subcluster. **A**: Jaccard, groups of genomes with similar accessory gene content. ClonaFrameML, logarithmic sum of recombining regions in the alignment of core genes. Subcluster, PopCOGenT subcluster number. **B**: Admixture blocks (ancestral recombination), patterns of shared ancestry within homologous regions across the core genome alignment. Blocks of the same color represent homologous regions shared by two lineages. The color of each block is assigned based on the lineage with the higher number of samples. FastGEAR detected three lineages: green (PopCOGenT subcluster 5.4), red (subclusters 5.0 + 5.3), and blue (subclusters 5.1 + 5.2). White blocks represent recent recombination, indicating putative recombination events occurring in some, but not all, members of a population.

### 3.5. Toxin genes

We focused primarily on key *Bcsl* virulence factors: (i) the *ces* operon, involved in cereulide production; (ii) insecticidal *Bt* toxin genes (*cry*, *cyt*, and *vip*); and (iii) the enterotoxin-encoding genes *nheABC, hblCDA, hlyII* and *cytK*.The *ces* operon was detected in 21 out of 294 samples, with a predominance in *B. paranthracis* subcluster 0.3 (Fig. 1, Supplementary Table S1). In this subcluster, 18 out of 36 genomes contained the operon. The *ces* operon was also identified in *B. paranthracis* subcluster 0.4 (2 out of 10 genomes) and in one *B. mycoides* genome.

The genes encoding insecticidal toxins (Cry, Cyt, and Vip) were detected in all samples within *Bt* subcluster 1.1, except for the four “non-*Bt*” genomes. These genes were also found in some samples of *Bt* cluster 9 (2/8 samples) and *Bt* cluster 10 (1/2). Additionally, they were detected in *Bcss* subcluster 1.0 (3/28), *B. wiedmannii* + “*B. pretiosus*” cluster 5 (3/24), *B. toyonensis* cluster 8 (2/9), and a single *B. albus* genome.

The *cytK-1* gene variant, specific to *B. cytotoxicus* (Fagerlung et al., 2004), was detected in all members of the corresponding cluster 2. The *cytK-2* variant was detected in 6 out of 14 PopCOGenT clusters, mainly occurred in *B. pacificus* subcluster 0.1 (4/4), *B. paranthracis* subcluster 0.4 (9/10), *B. paranthracis* subcluster 0.7 (2/2), *Bcss* + *Bt* cluster 1 (112/113), *B. anthracis* cluster 4 (8/11), *Bt* cluster 9 (4/8), and *Bt* cluster 10 (2/2). The *hlyII* gene was mainly found in *B. wiedmannii* + “*B. pretiosus*” cluster 5 (21/24), *B. anthracis* cluster 4 (9/11) and *Bt* cluster 9 (7/8).

The *hbl* operon predominantly occurred in *Bcss* + *Bt* cluster 1 (112/113), as well as in almost all members of *B. mycoides* cluster 6 (10/11), *B. toyonensis* cluster 8 (9/9), *Bt* cluster 9 (8/8), and *B. anthracis* cluster 11 (2/2). In *B. wiedmannii* + *“B. pretiosus”* cluster 5, the operon was present in all subclusters, except subcluster 5.4. Lastly, all samples, except 18SBCL773 (cluster 1), carried the *nhe* operon.

### 3.6. Antimicrobial and heavy metal resistance genes

Thereafter, we searched for the genes that might potentially be involved in antibiotic and heavy metal resistance in the *Bcsl* group. Using AMRFinderPlus, we detected beta-lactamase encoding genes (*bla1*, *bla2*, or *blaIII*) in all clusters, except for *B. cytotoxicus* cluster 2 and the unclustered genome 09CEB56 (Fig. 1).

The macrolide resistance genes, *mphL* and *mphM,* encoding macrolide 2’-phosphotransferase, were found in 17.0% (50/294) and 1.7% (5/294) of examined samples, respectively. The *mphL* gene was mainly detected in *B. paranthracis* subcluster 0.4 (8/10), *B. anthracis* cluster 4 (11/11), *B. luti* cluster 7 (3/3), *B. toyonensis* cluster 8 (9/9), and *Bt* cluster 9 (5/8). The *tet(45)* gene encoding a tetracycline efflux pump was found in 4.1% (12/294) of samples, while *tet(L)* was identified in one *B. anthracis* sample (10CEB21). The *vanR* and *vanS* genes, constituting the regulatory component of the *van* operon, were detected in 9.2% (27/294) of samples. These genes were annotated as the ‘vancomycin resistance response regulator transcription factor VanR’ and the ‘vancomycin resistance histidine kinase VanS,’ respectively. Additionally, we identified the *vanY* and *vanZ* genes, which represented the accessory components of the operon. The *vanY* gene was annotated as ‘VanY domain-containing protein,’ ‘LD-carboxypeptidase,’ or ‘D-alanyl-D-alanine carboxypeptidase’. Notably, the core resistance elements, encoded by the *vanHAX* gene cluster, were absent in all samples. Furthermore, we observed several structural variations (Fig. 5): *vanRSY, vanRSYZ, vanRS-lipoprotein-vanYZ*, and *vanRSYYZ*. In the genome 22SBCL0142_2 (*B. paranthracis* subcluster 0.4), we identified three variants of *vanRSY*. In the first variant, the genes were annotated as vancomycin resistance genes. In the second variant, *vanR* was annotated as ‘VanR-ABDEGLN family response regulator transcription factor’, and *vanS* as ‘vancomycin resistance histidine kinase VanS’, with a downstream carboxypeptidase was by the *dacC* gene. In the third variant, *vanR* was annotated as ‘DNA-binding response regulator OmpR family’, *vanS* as ‘Sensor histidine kinase WalK’, and *vanY* as ‘LD-carboxypeptidase LdcB LAS superfamily’.

**Figure 5.**
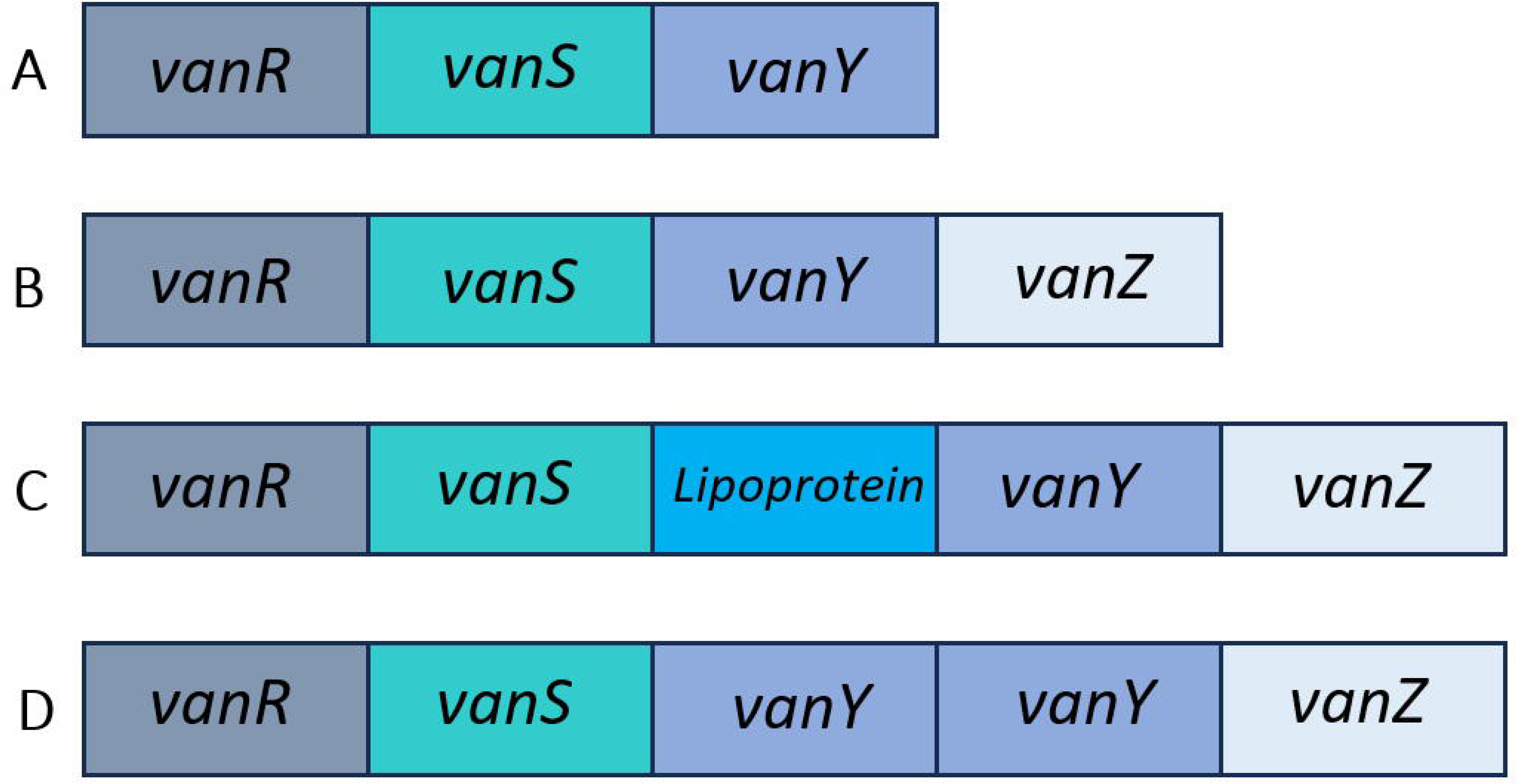
Structural variations in the gene cluster forming regulatory and accessory parts of the putative vancomycin resistance operon detected in *Bcsl.* **A**: *vanRSY* gene cluster. **B**: *vanRSYZ* gene cluster. **C**: *vanRSYZ* gene cluster including a lipoprotein-encoding gene. **D**: Gene cluster with duplicated *vanY*.

The mercury (*merB3R1EPAR2B2B1*) (Naguib et al., 2018) and arsenic (*arsRBCDA*) (Chien et al., 2019) resistance operons were observed in 3.1% (9/294) and 17.0% (50/294) of genomes, respectively (Fig. 1).

Finally, we reviewed the type of dishes associated with strains carrying antibiotic resistance genes (Supplementary Table S1). We noticed that the isolates carrying the *tet* genes were much more frequently associated with dishes marinated and/or containing spices (62% versus 7%) than the *tet*-negative isolates. Similarly, although with a smaller gap, we observed that the samples carrying *mphL* were more frequently associated with the “Fish, shellfish, and their products” FAO Level 1 category (16% versus 5%).

### 3.7. Antibiotic susceptibility phenotype

To confirm the phenotype of loss of antibiotic susceptibility due to the presence of antibiotic resistance genes (ARGs) previously highlighted, we carried out disk diffusion essays on 64 *Bcsl* isolates (Fig. 6, Supplementary Table S4). We first compared the susceptibility to ampicillin (a β-lactam class antibiotic) between *bla+* (n=53) and *bla-* (n=11) isolates. The isolates not carrying *bla* genes, predominantly belonging to *B. cytotoxicus* (10/11), demonstrated a large and significant increase in ampicillin susceptibility compared to *bla*+ isolates. Concerning erythromycin, a macrolide antibiotic, results revealed that the isolates carrying *mphL* (n=17) exhibited a significantly reduced inhibition diameter, compared to the *mphL*-isolates (n=47). According to the breakpoint defined by EUCAST (R<24 mm), 88% (15/17) of *mphL*+ isolates were resistant to macrolides. Similarly, *Bcsl* isolates carrying the *tet(45)* gene (n=12) exhibited a significantly reduced inhibition diameter when being exposed to tetracyclin (a cyclin class antibiotic), compared to the *tet(45)*-isolates (n=51). Moreover, a single isolate carrying the *tet(L)* gene showed a total lack of sensitivity to this antibiotic. Finally, the isolates carrying the *vanR* and *vanS* genes (n=20) showed a significant, albeit small, reduction in inhibition diameter than the isolates lacking these genes (n=44), after exposure to vancomycin, a glycopeptide-belonging antibiotic. However, according to EUCAST criteria (R<10 mm), all the isolates were susceptible to vancomycin.

**Figure 6.**
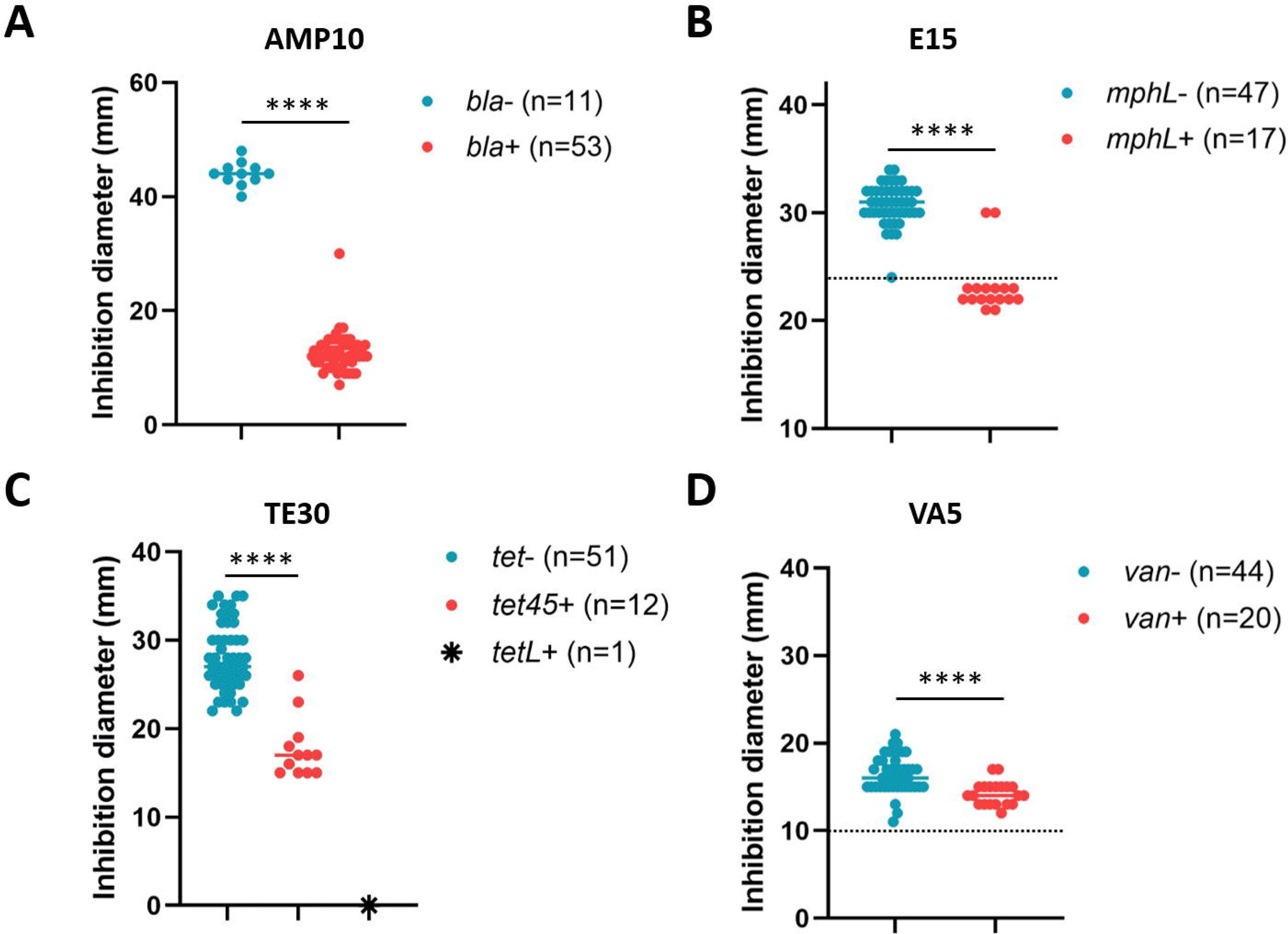
Antibiotic susceptibility testing of *Bcsl* performed using the Kirby–Bauer disk diffusion method (EUCAST method v13.0 and breakpoint tables for interpretation v15.0). For each antibiotic, isolates were selected based on the presence or absence of the identified antimicrobial resistance genes of interest. **A**: Resistance to ampicillin (10 µg, AMP10) examined in the *bla-* and *bla+* isolates; *bla*, beta-lactamase encoding genes. **B**: Resistance to erythromycin (15 µg, E15) resistance examined in the *mphL-* and *mphL+* isolates; *mphL*, macrolide 2’-phosphotransferase encoding gene. **C**: Resistance to tetracycline (30 µg, TE30) examined for the *tet-*, *tet(45)+* and *tet(L)+* isolates; *tet*, tetracycline efflux pump encoding gene. **D**: Resistance to vancomycin (5 µg, VA5) examined in the *van-* and *van+* isolates; *van*, putative regulatory part (*vanRS*) of the vancomycin resistance operon. For B and D, dotted lines indicate the respective values of EUCAST zone diameter breakpoints (Breakpoint Table v.14.0). ****, *p*-value<0.0001 (Mann-Whitney test). The figure was designed using GraphPad v.10.4.1.

### 3.8. Proportions of *Bcsl* populations

To search for associations between the most frequently found *Bcsl* populations and food categories, we applied a bootstrap-based approach to reflect the true proportions of the FBO-associated *Bcsl* populations in France. The most frequently detected populations were *Bcss* subcluster 1.0 (17.0±0.3%), *B. paranthracis* subcluster 0.3 (16.1±1.0%), and *Btk* subcluster 1.1 (7.6±1.0%) (Fig. 7). In FBOs where a single isolate was detected, the most frequent population was *B. paranthracis* subcluster 0.3, accounting for 6.3±1.4% of all FBO cases, followed by *Btk* (3.0±1.3%) and *Bcss* (2.8±0.5%).

**Figure 7.**
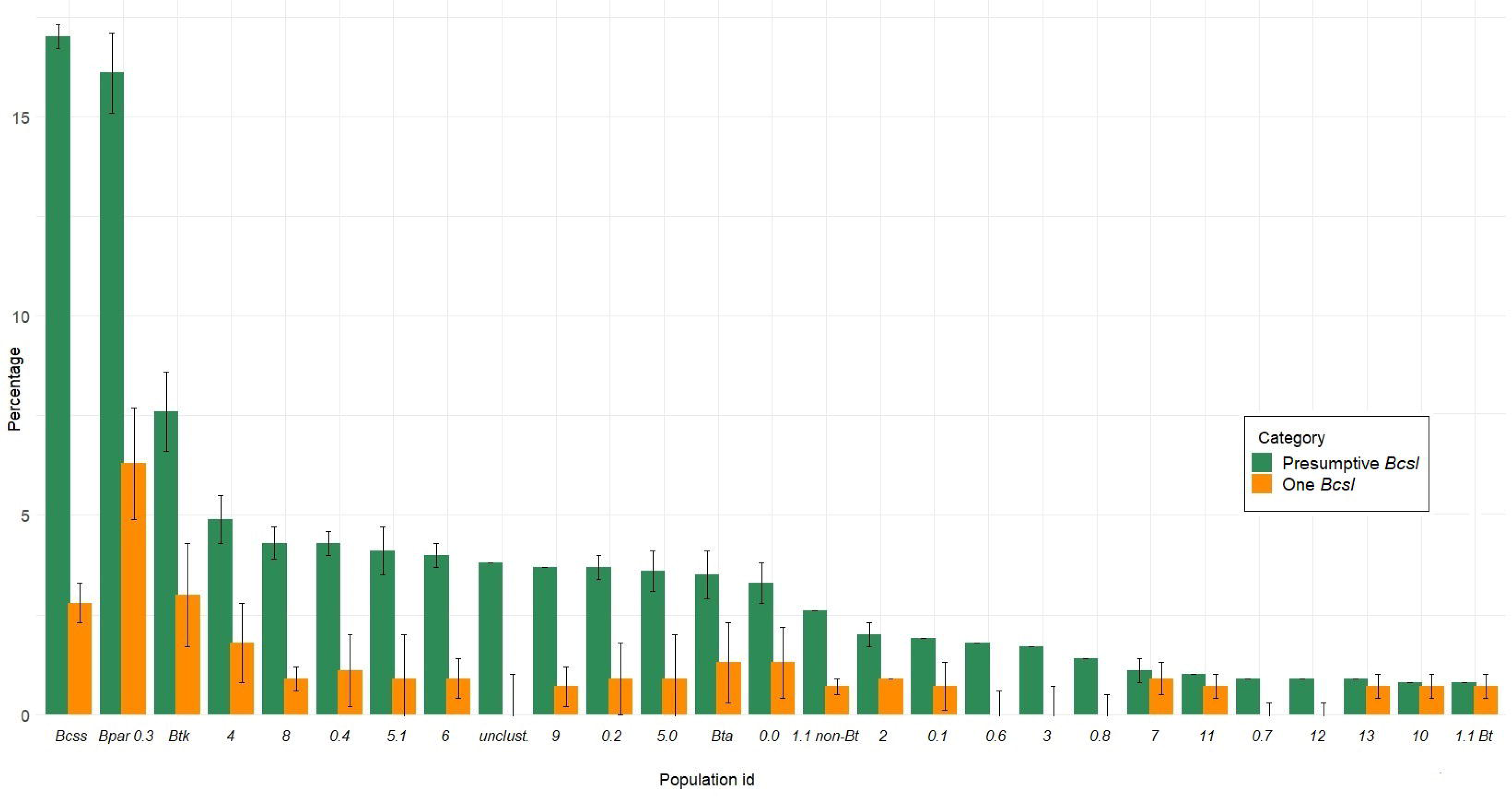
Proportions of *Bcsl* populations estimated from the collection of sequenced genomes sampled during foodborne outbreaks (FBO) in France between 2004 and 2023. **Presumptive *Bcsl***, the sample is counted even if another pathogen (another species or mix of *Bcsl* species) is identified in the FBO. **One *Bcsl***, only a single *Bcsl* isolate detected in the FBO. *Bcss*, *B. cereus sensu stricto* subcluster 1.0. *Bpar* 0.3, *B. paranthracis* subcluster 0.3. *Btk*, *Bt* subsp. *kurstaki* subcluster 1.1. 4, *B. anthracis* cluster 4. 8, *B. toyonensis* cluster 8. 0.4, *B. paranthracis* subcluster 0.4. 5.1, *B. wiedmannii + “B. pretiosus”* subclusters 5.1 + 5.2. 6, *B. mycoides* cluster 6. Unclust., unclustered genome. 9, *Bt* cluster 9. 0.2, *B. pacificus* subcluster 0.2. 5.0, *B. wiedmannii + “B. pretiosus”* subclusters 5.0 + 5.3 + 5.4. *Bta*, *Bt* subsp. *aizawai* subcluster 1.1. 0.0, *B. paranthracis* subclusters 0.0 + 0.5. 1.1 non-*Bt*, particular *Bt* strains from subcluster 1.1. 2, *B. cytotoxicus* cluster 2. 0.1, *B. pacificus* subcluster 0.1. 0.6, *B. paranthracis* subcluster 0.6. 3, *B. mobilis* cluster 3. 0.8, *B. tropicus* subcluster 0.8. 7, *B. luti* cluster 7. 11, *B. anthracis* cluster 11. 0.7, *B. paranthracis* subcluster 0.7. 12, *B. paramobilis* cluster 12; 13, *B. wiedmannii* cluster 13. 10, *Bt* cluster 10. 1.1 *Bt*, Bt ST13 from subcluster 1.1.

### 3.9. Distribution of *Bcsl* populations across food categories

Using previously generated random sample sets, we estimated the distribution of the entire *Bcsl* group and the three predominant populations across food categories. Statistically significant associations were found when using FoodEx2 classification (European Food Safety Authority, 2015) (Supplementary Table S1). At Level 1, the *Bcsl* group was most prevalent in the “Composite dishes” category (51±0.1%), followed by “Grains and grain-based products” (14±0.1%), and “Vegetables and vegetable products” (14±0.1%) (Fig. 8A). At Level 2, *Bcsl* was primarily found in “Dishes including ready-to-eat meals excluding soups and salads” (39±0.1%) and “Soups and salads” (15±0.1%), both subcategories of “Composite dishes”, followed by “Cereals and cereal primary derivatives” (13±0.1%), a subcategory of “Grains and grain-based products” (Fig. 8B). At Level 1, analysis of the three predominant populations revealed a significant positive association between *Bcss* and “Composite dishes” (*p*=0.029), where it was present in 73±3.9% of cases (Fig. 8A). *B. paranthracis* subcluster 0.3 was detected in “Grains and grain-based products” and “Starchy roots and tubers” in 26±5.8% and 10±3.7% of cases, respectively, although these associations were not statistically significant when comparing with the overall *Bcsl* background. Similarly, *Btk* was found in “Fruit and fruit products” in 13±6.4% of cases, but without statistical significance. At Level 2, *B. paranthracis* subcluster 0.3 showed a significant positive association with “Cereals and cereal primary derivatives” (*p*=0.033), accounting for 24±5.5% of cases. Additionally, *Btk* was significantly associated with “Soups and salads” (34±12.2%) (*p*<0.0001), which primarily included raw vegetable-based salads (Fig. 8B, Supplementary Fig. S1).

**Figure 8.**
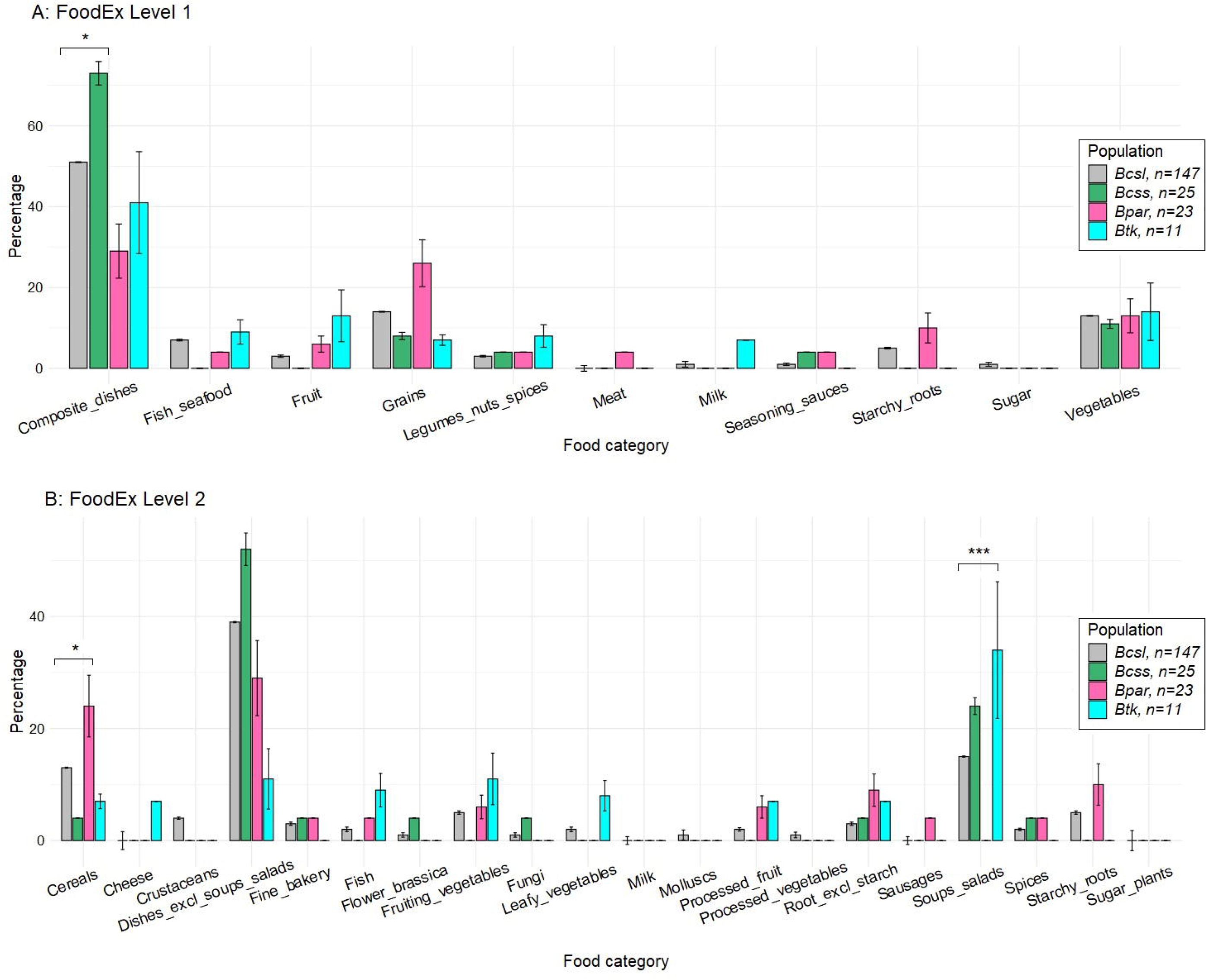
Distribution of the entire *Bcsl* group and the predominant populations across different food categories determined using FoodEx2 exposure classification Level 1 and 2 (European Food Safety Authority, 2015). **A: FoodEx Level 1**. Composite_dishes, “Composite dishes” category. Fish_seafood, “Fish, seafood, amphibians, reptiles and invertebrates”. Fruit, “Fruit and fruit products”. Grains, “Grains and grain-based products”. Legumes_nuts_spices, “Legumes, nuts, oilseeds and spices”. Meat, “Meat and meat products”. Milk, “Milk and dairy products”. Seasoning_sauces, “seasoning sauces and condiments”. Starchy_roots, “Starchy roots or tubers and products thereof sugar plants”. Sugar, “Sugar and similar confectionery and water-based sweet desserts”. Vegetables, “Vegetables and their products”. **B: FoodEx Level 2**. Cereals, “Cereals and cereal primary derivatives”. Cheese, “Cheese”. Crustaceans, “Crustaceans”. Dishes_excl_soups_salads, “Dishes including ready-to-eat meals excluding soups and salads”. Fine_bakery, “Fine bakery wares”. Fish, “Fish meat”. Flower_brassica, “Flowering brassica”. Fungi, “Fungi, mosses and lichens”. Leafy_vegetables, “Leafy vegetables”. Milk, “Milk and dairy powders and concentrates”. Mollusks, “Mollusks”. Processed_fruit, “Processed fruit products”. Processed_vegetables, “Processed or preserved vegetables and similar”. Root_excl_starch, “Root and tuber vegetables excluding starchy and sugar”. Sausages, “Sausages”. Soups_salads, “Soups and salads”. Spices, “Spices”. Starchy_roots, “Starchy roots and tubers”. Sugar_plants, “Sugar plants”. *Bcsl,* the *B. cereus sensu lato* entire group. *Bcss, B. cereus sensu stricto* subcluster 1.0. *Bpar, B. paranthracis* subcluster 0.3. *Btk, Bt* subsp. *kurstaki* subcluster 1.1.

## 4. DISCUSSION

In this study, we resolved the population structure of *Bcsl* collected during FBOs in France between 2004 and 2023.

First, when examining virulence genes, we observed putative signatures of horizontal transfer, notably for the *cry/cyt/vip* genes and the *ces* operon. For instance, although plasmid-associated genes encoding insecticidal toxins (Cry, Cyt, and Vip) are characteristic of *Bt* (Schnepf et al., 1998), we also identified them in the other species, comprising three cases of *Bcss*. Consistently, other *Bcsl* species possessing insecticidal toxin genes were described in previous studies (Lazarte et al., 2018; Sauka et al., 2022). Furthermore, Sauka et al. (2022) demonstrated that the *B. toyonensis* strain Bto-UNVM_94, which carries a *cry* gene, can produce parasporal crystals with insecticidal activity. Similarly, the plasmid-located *ces* operon (Ehling-Schulz et al., 2006), encoding genes involved in the production of the emetic toxin, is mostly found in a part of *B. paranthracis* ST26 samples, as also revealed in previous studies (Caroll et al., 2022b; Frentzel et al., 2024). Furthermore, the absence of the *ces* operon in the other *B. paranthracis* ST26 strains indicates the process of gene gain and loss within the entire ST26 clade, as previously detailed by Caroll and Wiedmann (2020). We also revealed one case of *B. mycoides* carrying the *ces* operon, suggesting that *Bcsl*, not classified within *panC* group III, may acquire the genes responsible for cereulide production. Indeed, several *B. mycoides* strains isolated from soil were shown to carry the *cesHPTABCD* operon as part of the composite transposon Tn*ces* (Mei et al., 2014). Also, Frentzel et al. (2024) described an emetic *B. mobilis* strain isolated from meat salad, able to produce cereulide. In the study of Pheepakpraw et al. (2023), the presence of the *cesA* and *cesB* gene fragments was confirmed for several FBO- and clinic-associated strains, including *Bt* ST8, *Bt* ST15, and *Bcsl* ST2804 and ST2805. For the other virulence factors, we demonstrated that *nhe* was present in almost all samples, while *hbl, hlyII* and *cytK-2* were found in various lineages. This supports the findings of Böhm et al. (2015), who showed that the *nhe* operon was present in all *Bcsl* and was vertically inherited with extremely rare cases of gene loss. In contrast, *hbl* and *cytK-2* were shown to be subject to horizontal transfer (Böhm et al., 2015). Finally, we observed the predominant occurrence of the *hbl* operon in *panC* phylogenetic groups II, IV, V, and VI. This aligns with the findings of Carroll et al. (2017), who demonstrated a statistically significant association between the presence of the operon and these clades.

Furthermore, we sporadically identified the antimicrobial and heavy metal resistance genes, such as *tet(45)*, *tet(L)*, *mphL*, *ars*, and *mer*, within distinct *Bcsl* populations, also supposing their acquisition by horizontal transfer and suggesting the adaptation of *Bcsl* to specific environmental conditions. We noticed that the *Bcsl* isolates carrying the *tet* genes were much more frequently detected in dishes containing spices. Given that most spices in France are imported, we speculate that this may reflect a potential spread of ARG-carrying *Bcsl* across geographical borders, particularly from regions with higher antibiotic pressure (Okeke and Edelman, 2001). Similarly, the *mph* genes were frequently associated with seafood products, for which the spread of ARGs is a well-documented concern, both in marine environments (Ferraro et al., 2024) and fish farming areas (Helsens et al., 2020).

By contrast, we found that all *Bcsl*, except *B. cytotoxicus* cluster 2, possessed genes encoding various beta-lactamases. Indeed, numerous studies demonstrated 98-100% resistance of *Bcsl* isolates to penicillin, cefotaxime, ampicillin and oxacillin, concluding natural resistance of *Bcsl* to beta-lactams (Glasset et al., 2018; Bianco et al., 2021; Jung et al., 2022; Rajalingam et al., 2022; Mohammadi et al., 2023). To date, this is the first time that a specific *Bcsl* population (*B. cytotoxicus*) is exhaustively associated with the absence of beta-lactamases and shows a drastically increased sensitivity phenotype to a beta-lactam antibiotic, compared to the rest of the group.

Thereafter, when searching for the *van* operon, we detected several variants of the *vanRSYZ* gene cluster with no resistance-associated genes (*vanH* encoding dehydrogenase, *vanA/B/D/F/M* encoding D-ala-D-lactate ligase, and *vanX* encoding D,D-dipeptidase) (Stogios and Savchenko, 2020). Indeed, the phylogenetic trees reconstructed by Kardos et al. (2024) for the vancomycin resistance genes in *Bacilli* and *Clostridia* sequences available in the NCBI database showed that the acquisition of *vanRS*, with or without *vanYZ*, could occur in several clades. These acquisitions may either be associated with resistance-related genes or exist independently, suggesting that *vanRS*-like regulatory systems could potentially regulate other genes unrelated to the *van* operon (Kardos et al., 2024). This is also supported by numerous studies showing 95-100% susceptibility of *Bcsl* strains to vancomycin (Glasset et al., 2018; Bianco et al., 2021; Jung et al., 2022; Rajalingam et al., 2022; Mohammadi et al., 2023).

To sum up, future research should complement gene presence-absence analysis with gene expression analysis, genetic and structural variation analysis, and genome-wide association studies to link specific genotypes with phenotypic traits, such as toxin production levels and antibiotic resistance. This is crucial for the *Bcsl* group in the context of public health, as non-pathogenic strains may acquire virulence factors and develop antibiotic resistance via horizontal gene transfer and mutation.

The clustering of *Bcsl* strains enabled us to detect three composite PopCOGenT clusters, each representing a group of closely related species: *B. paranthracis* + *B. pacificus* + *B. tropicus* cluster 0, *Bcss + Bt* cluster 1, and *B. wiedmannii* + *“B. pretiosus”* cluster 5. In these clusters, FastGEAR identified numerous admixture blocks within lineages, suggesting either shared ancestry or recombination. Metrics from ClonalFrameML show that recombination has a greater overall impact than mutation in the examined *Bcsl* clusters, representing an intermediate level between clonal and true recombining bacterial species (Vos and Didelot, 2008; Patiño-Navarrete and Sanchis, 2017). Our estimations are consistent with the findings of Didelot and Falush (2007), who estimated r/m at 1.5-2.8 using the MLST data of *Bcsl.* Thus, we conclude that recombination may serve as an important mechanism for *Bcsl* to share fitness-enhancing alleles.

The smallest *B. wiedmannii* + *“B. pretiosus”* composite cluster 5 comprises two species. The *B. wiedmannii* type strain was originally isolated from raw milk and characterized as a psychrotolerant isolate producing hemolysin BL and non-hemolytic enterotoxin, as well as showing cytotoxicity to HeLa cells (Miller et al., 2016). *“B. pretiosus”*, effectively published as species phylogenetically close to *B. wiedmannii*, was initially isolated from the rhizosphere of alfalfa (Robas Mora et al., 2023). The significant Groups of samples detected based on the accessory genome did not entirely align with the population structure. This suggests that the *B. wiedmanii* + *“B. pretiosus”* cluster occupies a broad ecological niche, with strains adapting to local environmental conditions through a unique set of accessory genes.

The second *Bcss* + *Bt* cluster 1, consists of two subclusters, each representing a distinct species: *Bt* and *Bcss*. Within *Bt* subcluster 1.1, the application of Bt_typing (Fichant et al., 2022) allowed us to identify several *Bt* lineages: *Bta*, *Btk*, *Bt* ST13, and a group of samples lacking *Bt*-specific genetic markers as well as *cry*, *cyt*, or *vip* genes, suggesting that they may represent particular *Bt* strains. In addition, we want to emphasize that the study by Fichant et al. (2022) aimed to develop the workflow specifically for pesticide and FBO-associated *Bta* and *Btk* strains rather than for the full diversity of *Bt*. The *Bcss* subcluster 1.0 is represented by a single PopCOGenT/fastGEAR population showing complex population structure. The subcluster includes various STs, with predominant ST24. Furthermore, clinical *Bcss* samples are phylogenetically closely related to the various FBO-associated strains, reinforcing the ability of *Bcss* to disseminate among diverse environments. This also suggests that these *Bcss* lineages can cause both FBOs and extra-intestinal infections, highlighting the potential of a digestive tract as an entry for systemic infections.

The analysis of accessory gene content in *Bcss* ST24 and *Bta* revealed that each lineage was enriched with a specific set of genes associated with the “Mobilome: phages, transposons” functional category. Indeed, the mobilome of *Bcsl* includes various transposable elements (TEs), such as insertion sequences and class II transposons that are often associated with virulence genes (Mei et al., 2014; Fayad et al., 2019). Fayad et al. (2019) found several insertion sequences (IS231, IS232, and IS240) to be structurally associated with delta-endotoxin-encoding genes of *Bt*. This highlights the value of further study to search for associations between TEs and accessory genes in our *Bscl* genomic dataset, which requires complete genome assemblies for a detailed investigation of gene cassettes.

Finally, we found that *Bt* subcluster 1.1 was phylogenetically closer to the *Bcss* subcluster 1.0 than to *Bt* from the distinct clusters 9 and 10, thereby suggesting that *Bt* might represent a polyphyletic group. The *Bt* from clusters 9 and 10 represent various *Bt* sequence types, including *Bt* serovar *israelensis* (ST16 and ST197) and *Bt* serovar *morrisoni* (ST23) (Supplementary Table S1) (Wang et al., 2018). Notably, only three out of ten strains carry delta-endotoxin-encoding genes. This observation suggests the sporadic loss or acquisition of insecticidal toxin-encoding genes within these *Bt* phylogenetic groups.

Therefore, genetic and phenotypic overlap within the *Bcss*-*Bt* species complex may complicate the differentiation between environmental *Bt* strains and pesticidal strains, as well as *Bt* and *Bcss* strains carrying *cry* genes, using laboratory methods such as PCR of the *panC* gene or microscopic observation of insecticidal crystals. An accurate identification of pesticidal *Bt* remains crucial, given the increasing evidence that *Bt* contaminates food products (EFSA Panel on Biological Hazards, 2016; Frentzel et al., 2014; Caroll et al., 2022b), disseminates in hospital environments (Muigg et al., 2022), and is associated with FBOs (Bonis et al., 2021; Biggel et al., 2022a, 2022b). Nevertheless, the initial steps to detect pesticidal *Bt* strains, primarily derived from *Bta* and *Btk* subspecies, were performed in by Frentzel et al. (2020), Bonis et al. (2021), Biggel et al. (2022a), and Fichant et al. (2022). Frentzel et al. (2020) found that some *Bt* strains isolated from vegetables differed from pesticidal strains by ≤4 wgSNPs. Biggel et al. (2022a) demonstrated that the genetic distance between several food- and FBO-associated *Bta* and *Btk* strains and pesticidal strains was no more than 2 cgSNPs or 6 wgSNPs, thereby suggesting a pesticidal origin for the food- and FBO-associated strains. Bonis et al. (2021) showed that pesticidal and FBO-associated *Bt* strains formed four phylogenetic clusters (a to d), with the median cgSNP distances ranged from 0 to 2 within each cluster. However, to date, there is no accepted threshold of genetic distance between pesticidal and non-pesticidal *Bt* genomes to classify them as the same strain. Subsequently, Fichant et al. (2022) proposed the approach to detect accessory genes specific for pesticidal and FBO-associated *Bt* strains at the species level (*Bt* and non-*Bt*), subspecies level (*Bta* and *Btk*), and cluster level (a to d). Further precise detection of *Bt* strains remains a subject for future work, requiring the identification of finer genetic markers such as SNPs or indels. This is crucial for *Bt* strain-specific risk assessments.

The largest cluster, *B. paranthracis* + *B. pacificus* + *B. tropicus* cluster 0, comprises three closely related species. Among them, *B. paranthracis* subcluster 0.3 contains the highest number of samples and consists of several strains, with predominant ST26 and ST266. Complex population structure of this cluster suggests a broad ecological niche, allowing the populations to adapt to a wide range of environments. The three species were initially isolated from ocean sediment (Liu et al., 2017). *B. paranthracis* is often associated with foodborne outbreaks (Messelhäusser et al., 2014). *B. pacificus* is found in soil, with some strains showing plant-growth promoting properties (Ma et al., 2024). *B. tropicus* is also found in soil and sludge environments (Thakur et al., 2021; Shen et al., 2022). Recombination likely plays a significant role in shaping the population structure of this cluster, especially since the R/θ and r/m ratios were higher compared to those for clusters 1 and 5. We revealed several strains, such as 11CEB28 (*B. tropicus* subcluster 0.8), 18SBCL174 (*B. paranthracis* subcluster 0.7) and 22SBCL0891_1 (*B. paranthracis* subcluster 0.7), that demonstrated the highest number of recent recombination blocks and the longest recombining regions, suggesting they are products of recent recombination between distinct populations or have recently inherited these signatures.

The analysis of accessory gene content revealed two distinct groups, Group 33 and Group 41, within *B. paranthracis* subcluster 0.3, each possessing specific accessory genes likely suited to specific environmental conditions. Interestingly, the accessory genes of *B. paranthracis* subclusters 0.0 and 0.5, and the clinical samples GCF_021014465, GCA_029696605, and GCF_036851755, are enriched in ‘Cell wall/membrane/envelope biogenesis’ COG functional category. These genes may play a role in the synthesis of cell wall components, including virulence factors. These molecules may contribute to host-pathogen interactions, such as colonization of host tissues and modulation of the host immune system (King and Roberts, 2016). This is particularly crucial for both FBO-related and clinical *Bcsl* strains.

After characterizing the *Bcsl* populations, we identified that the most frequent FBO-related groups were *Bcss*, *B. paranthracis* subcluster 0.3 including emetic strains, and *Btk.* The predominance of the commonly known foodborne pathogens *Bcss* and *B. paranthracis* was also demonstrated in other studies (Messelhäusser et al., 2014; Frentzel et al., 2024). The predominance of *Btk* can be explained by the widespread use of *Bt* pesticide strains and subsequent contamination of food, particularly vegetable-based meals, which aligns with the findings of Bonis et al. (2021). Furthermore, *B. paranthracis* subcluster 0.3 showed significant positive association with starchy foods, such as cereals, while *Bcss* was associated with composite dishes. These findings align with the study by Messelhäusser et al. (2014), who reported that cereulide-producing strains isolated during FBOs in Bavaria between 2007 and 2013 predominantly occurred in cereals. Additionally, various reports (Osimani et al., 2018; Santé Publique France, 2024) highlighted that the *Bcsl* group was commonly linked to composite dishes, including pasta- and rice-based dishes. This indicates the elevated risks associated with these food categories and highlights the challenge of tracing the initial source of contamination.

## 5. CONCLUSION

In our study, we analyzed the population structure of a large collection of FBO-associated *Bcsl* strains collected in France between 2004 and 2023. Using 294 genomes from 183 FBOs, we identified 14 PopCOGenT clusters corresponding to validly and effectively published species. Three of these clusters exhibited a composite structure, each composed of several distinct populations. The predominant FBO-associated populations in France are *Bcss* subcluster 1.0 comprising various STs, *B. paranthracis* subcluster 0.3 including emetic strains, and *Btk* subcluster 1.1. In the perspective, the identified populations will be used for further investigations, in particular to search for correlations with epidemiological indicators, such as the percentage of affected individuals, age, symptoms, infectious dose, and collection point. Furthermore, we found that several FBO-associated *Bcsl* strains belonged to the same phylogenetic clades as clinical strains, highlighting the need for antibiotic susceptibility assessments. Notably, we found that *B. cytotoxicus,* which lacks beta-lactamase-encoding genes, was significantly more sensitive to ampicillin than other *Bcsl* lineages, which are considered to be naturally resistant to beta-lactams. Additionally, the strains carrying macrolide- and tetracycline resistance genes showed reduced susceptibility to erythromycin and tetracycline. Finally, we identified recombining *Bcsl* strains and putative signatures of horizontal transfer of the *Bcsl* toxin-encoding genes between distinct lineages. This is crucial, as pathogenic strains may transfer virulence factors and fitness-enhancing alleles to non-pathogenic strains. These findings should be considered in the surveillance of foodborne outbreaks associated with the *B. cereus* group.

## AUTHOR CONTRIBUTIONS

KM: Conceptualization, Investigation, Formal analysis, Visualization, Writing – original draft. SP: Methodology, Investigation. OF: Conceptualization, Supervision, Validation, Writing – review and editing. MB: Conceptualization, Investigation, Funding acquisition, Supervision, Validation, Writing – review and editing.

## ACKNOWLEGEMENTS

We thank all the district veterinary and food analysis laboratories for carrying out *Bcsl* detection and transmitting isolates together with epidemiologic data to the Food Safety Laboratory. We also thank Laurent Guillier for the fruitful discussions on the project.

## DATA AVAILABILITY STATEMENT

The accessions for the raw reads of the studied *Bcsl* strains are provided in Supplementary Table S1.

