## Supplementary figures and images for "Population structure of *Bacillus cereus sensu lato* associated with foodborne outbreaks in France between 2004 and 2023"

### Fig. S1

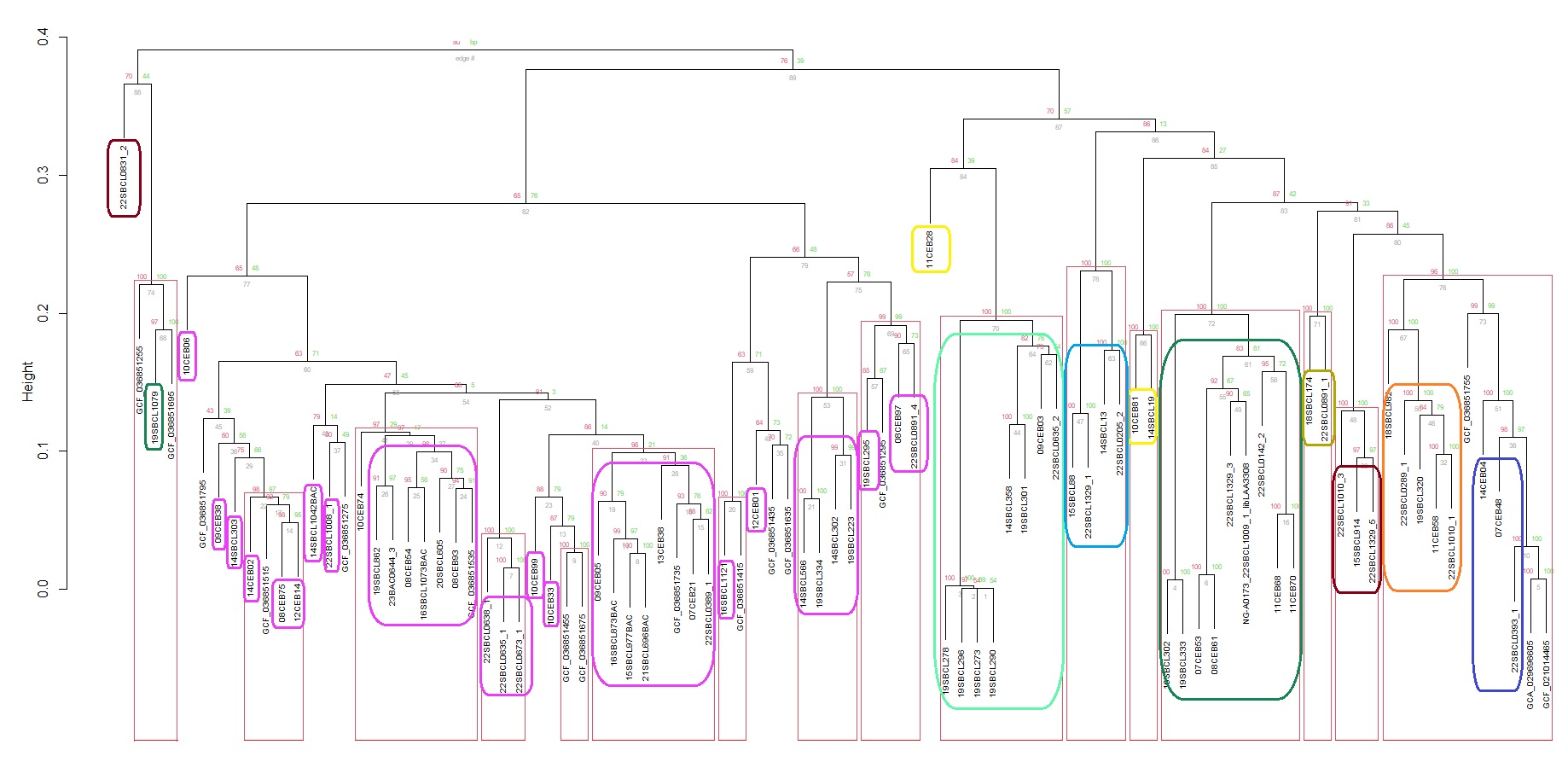

### Fig. S2

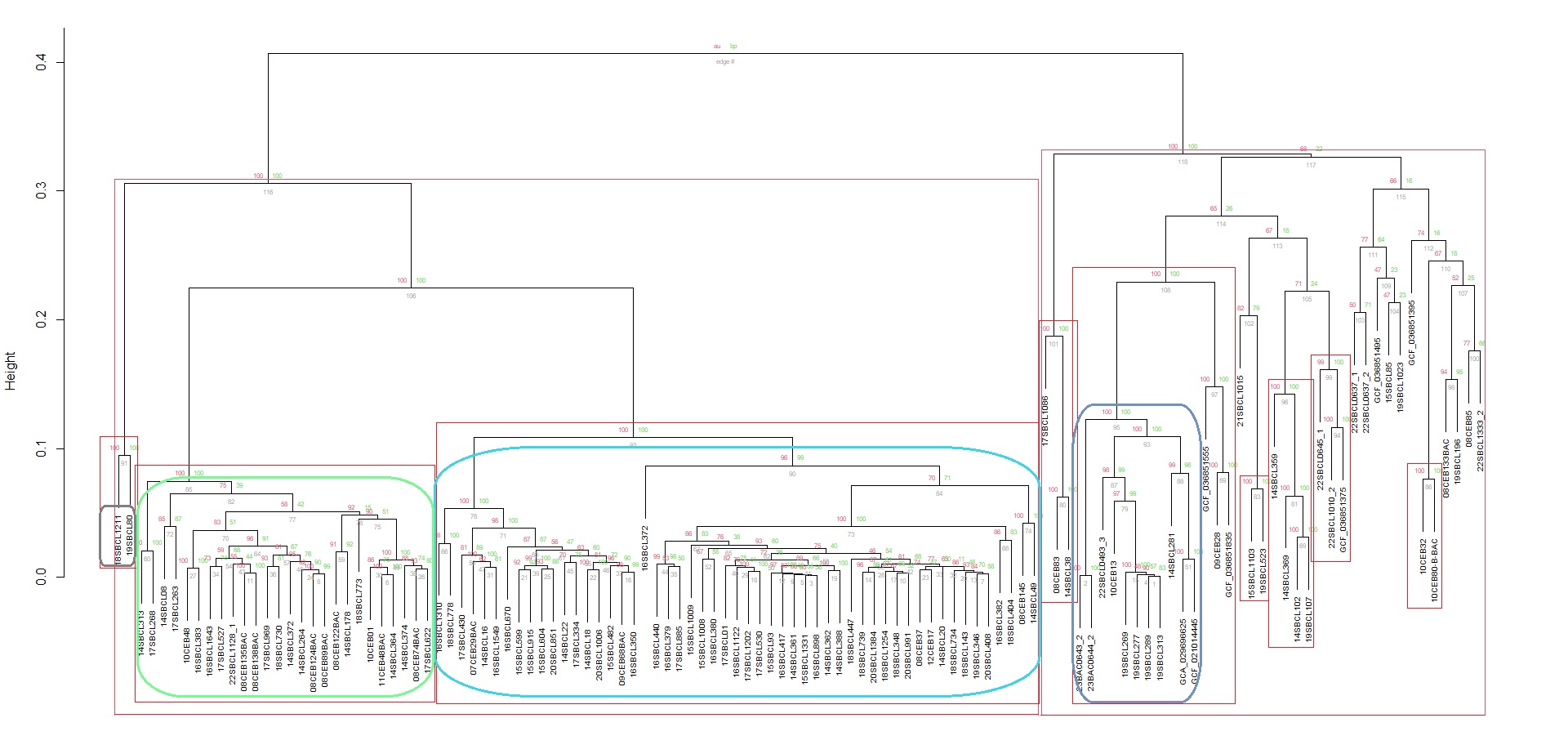

### Fig. S3

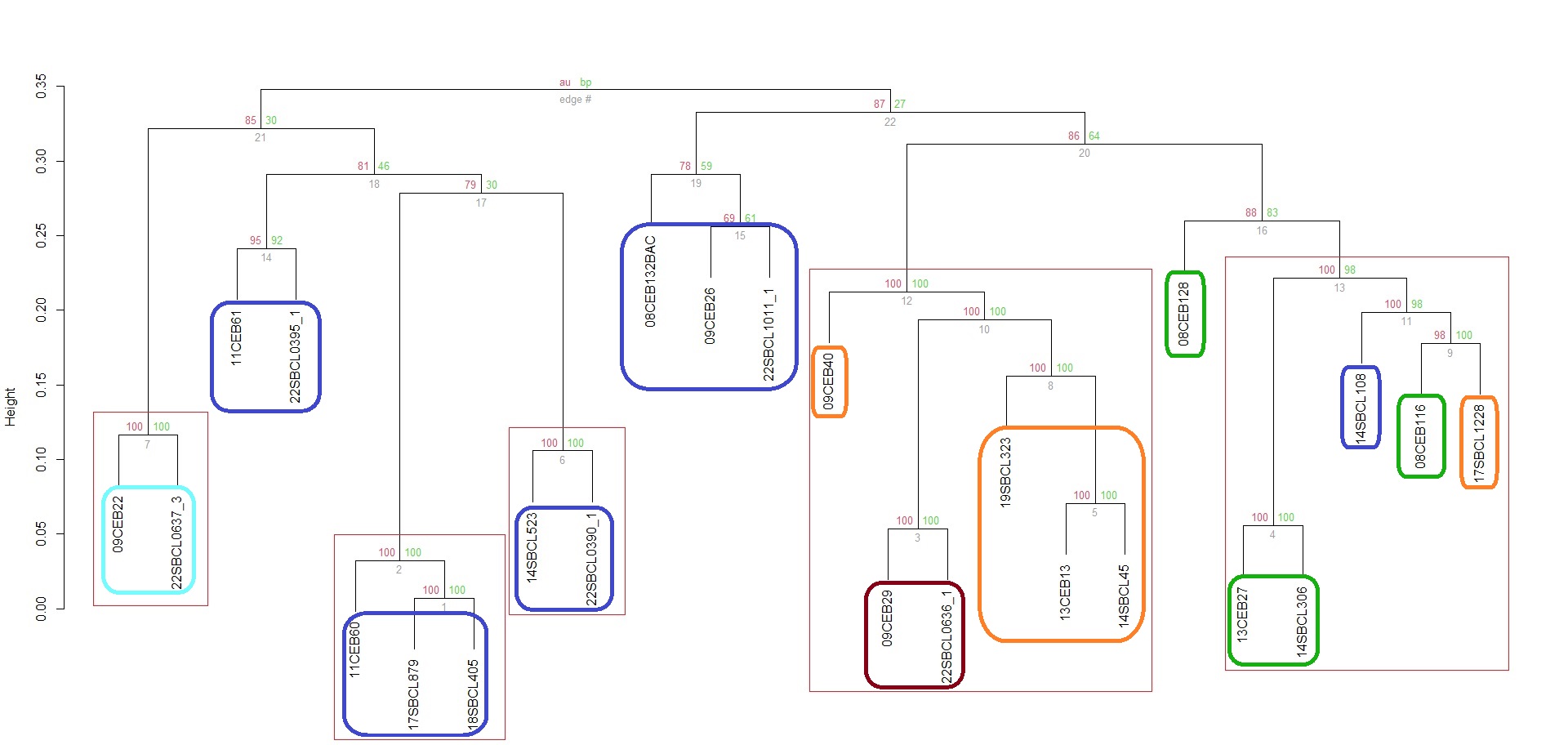
