## Supplementary material for "Population structure of *Bacillus cereus sensu lato* associated with foodborne outbreaks in France between 2004 and 2023": Fig. S1, S2, S3

**SUPPLEMENTARY FIGURES**

**Figure S1**. Groups of genomes in *B. paranthracis* + *B. pacificus* + *B. tropicus* PopCOGenT cluster 0 showing similar gene content. The groups were identified by calculating pairwise Jaccard indices with the pvclust R package v. 2.2.0 (Suzuki et al., 2019). Grey values, group numbers. Red values, approximately unbiased p-values. Green values, bootstrap probability values. Height, dissimilarity measure at which groups are merged in hierarchical clustering. Significant results are outlined in red. Populations resolved within the PopCOGenT cluster: *B. paranthracis* subcluster 0.0 (orange), *B. pacificus* subcluster 0.1 (light blue), *B. pacificus* subcluster 0.2 (light green), *B. paranthracis* subcluster 0.3 (purple), *B. paranthracis* subcluster 0.4 (dark green), *B. paranthracis* subcluster 0.5 (blue), *B. paranthracis* subcluster 0.6 (brown), *B. paranthracis* subcluster 0.7 (khaki), *B. tropicus* subcluster 0.8 (yellow). The GCA* and GCF* sample ids correspond to the clinical *B. paranthracis* samples (**Supplementary Table S2**).

**Figure S2**. Groups of genomes in *Bcss* + *Bt* PopCOGenT cluster 1 showing similar gene content. The groups were identified by calculating pairwise Jaccard indices with the pvclust R package v. 2.2.0 (Suzuki et al., 2019). Grey values, group numbers. Red values, approximately unbiased p-values. Green values, bootstrap probability values. Height, dissimilarity measure at which groups are merged in hierarchical clustering. Significant results are outlined in red. Populations resolved within the PopCOGenT cluster: subcluster 1.0 (*B. cereus sensu stricto* ST24 outlined in blue and other samples forming Group 118), subcluster 1.1 (*B. thuringiensis* subsp. *aizawai* outlined in green, *B. thuringiensis* subsp. *kurstaki* outlined in cyan, and *B. thuringiensis* ST13 outlined in grey). The samples 15SBCL85, 19SBCL1023, 22SBCL0637_1, and 22SBCL0637_2 belong to subcluster 1.1 but are grouped together with *Bc* into Group 18. The GCA* and GCF* sample ids correspond to the clinical *Bc* samples (**Supplementary Table S2**).

**Figure S3**. Groups of genomes in *B. wiedmannii* + *‘’B. pretiosus’’* PopCOGenT cluster 5 showing similar gene content. The groups were identified by calculating pairwise Jaccard indices with the pvclust R package v. 2.2.0 (Suzuki et al., 2019). Grey values, group numbers. Red values, approximately unbiased p-values. Green values, bootstrap probability values. Height, dissimilarity measure at which groups are merged in hierarchical clustering. Significant results are outlined in red. Populations resolved within the PopCOGenT cluster: subcluster 5.0 (brown), subcluster 5.1 (blue), subcluster 5.2 (cyan), subcluster 5.3 (orange), subcluster 5.4 (green).
