## Supplementary material for "Population structure of *Bacillus cereus sensu lato* associated with foodborne outbreaks in France between 2004 and 2023": Table S2

**Table S2**. *B. cereus sensu stricto* and *B. paranthracis* clinical strains used in the study.

| **Accession** | **Genomospecies** | **Closest type strain** | **ST** |
| --- | --- | --- | --- |
| GCA_029696625 | *B. cereus sensu stricto* | *B. cereus sensu stricto* | 24 |
| GCF_021014445 | *B. cereus sensu stricto* | *B. cereus sensu stricto* | 24 |
| GCF_036851555 | *B. cereus sensu stricto* | *B. cereus sensu stricto* | 73 |
| GCF_036851835 | *B. cereus sensu stricto* | *B. cereus sensu stricto* | 73 |
| GCF_036851395 | *B. cereus sensu stricto* | *B. cereus sensu stricto* | 1916 |
| GCF_036851375 | *B. cereus sensu stricto* | *B. cereus sensu stricto* | 1969 |
| GCF_036851495 | *B. cereus sensu stricto* | *B. cereus sensu stricto* | 2779 |
| GCF_036851275 | *B. mosaicus* | *B. paranthracis* | 26 |
| GCF_036851435 | *B. mosaicus* | *B. paranthracis* | 26 |
| GCF_036851455 | *B. mosaicus* | *B. paranthracis* | 26 |
| GCF_036851515 | *B. mosaicus* | *B. paranthracis* | 26 |
| GCF_036851535 | *B. mosaicus* | *B. paranthracis* | 26 |
| GCF_036851635 | *B. mosaicus* | *B. paranthracis* | 26 |
| GCF_036851675 | *B. mosaicus* | *B. paranthracis* | 26 |
| GCF_036851735 | *B. mosaicus* | *B. paranthracis* | 26 |
| GCF_036851795 | *B. mosaicus* | *B. paranthracis* | 26 |
| GCF_036851295 | *B. mosaicus* | *B. paranthracis* | 144 |
| GCF_036851415 | *B. mosaicus* | *B. paranthracis* | 164 |
| GCA_029696605 | *B. mosaicus* | *B. paranthracis* | 611 |
| GCF_021014465 | *B. mosaicus* | *B. paranthracis* | 611 |
| GCF_036851255 | *B. mosaicus* | *B. paranthracis* | 770 |
| GCF_036851695 | *B. mosaicus* | *B. paranthracis* | 770 |
| GCF_036851755 | *B. mosaicus* | *B. paranthracis* | 994 |
