## Supplementary material for "Population structure of *Bacillus cereus sensu lato* associated with foodborne outbreaks in France between 2004 and 2023": Table S3

**Table S3**. Number of group-specific genes identified for PopCOGenT clusters, PopCOGenT subclusters, and Jaccard groups

| Cluster | Subcluster / Group Jaccard index | Number of group-specific genes* | Number of COG ids | Significant COGs** |
| --- | --- | --- | --- | --- |
| *B. wiedemanni +*  *“B. pretiosus”* | *B. wiedemanni +*  *“B. pretiosus”* | 56 (40) | 49 (37) |  |
|  | Group 12  subcluster  5.0 + 5.3 | 155 (59) | 83 (33) |  |
|  | Group 13 | 0 (0) | 0 (0) |  |
| *Bcss + Bt* | *Bcss* | 52 (34) | 19 (17) |  |
|  | *Btk* | 1208 (1168) | 520 (260) |  |
|  | *Bta* | 611 (562) | 223 (109) | X: Mobilome: prophages, transposons |
|  | Group 95  *Bcss* ST24 | 216 (216) | 101 (75) | X: Mobilome: prophages, transposons |
| *B. paranthracis +*  *B. pacificus +*  *B. tropicus* | *B. paranthracis* | 138 (137) | 106 (79) |  |
|  | *B. pacificus* | 60 (55) | 45 (39) |  |
|  | Group 33  *B. paranthracis*  subcluster 0.3 ST26 | 37 (37) | 16 (14) |  |
|  | Group 41  *B. paranthracis*  subcluster 0.3 ST26 | 0 (0) | 0 (0) |  |
|  | Group 70  *B. pacificus*  subcluster 0.2 | 272 (235) | 162 (115) |  |
|  | Group 72  *B. paranthracis*  subcluster 0.4 | 148 (86) | 91 (70) |  |
|  | Group 76  *B. paranthracis*  subcluster 0.0 + 0.5 | 120 (91) | 85 (64) | M: Cell wall/membrane/envelope biogenesis |
| *B. anthracis*  cluster 4 | *B. anthracis* cluster 4 | 70 (47) | 56 (44) |  |
| *B. cytotoxicus*  cluster 2 | *B. cytotoxicus*  cluster 2 | 1810 (1794) | 1532 (745) |  |
| *B. mycoides*  cluster 6 | *B. mycoides* cluster 6 | 303 (218) | 228 (148) |  |
| *B. thuringiensis* cluster 9 | *B. thuringiensis* cluster 9 | 195 (142) | 121 (88) | K: Transcription |
| *B. toyonensis*  cluster 5 | *B. toyonensis*  cluster 5 | 341 (245) | 280 (163) | K: Transcription |

*Group-specific genes were identified using Scoary v. 1.6.16 (Brynildsrud et al., 2016) with 80% specificity, 80% sensitivity, and Benjamini-Hochberg correction. In parentheses: number of genes identified at 100% specificity. **Significant COGs were identified using MicrobiomeProfiler (Wu et al., 2021). In parentheses: number of unique COGs.
